# Temporal patterns of dominance in a hummingbird assemblage

**DOI:** 10.64898/2025.12.05.692456

**Authors:** Nicolas Tellez-Colmenares, Ana Melisa Fernandes, F. Gary Stiles, Alejandro Rico-Guevara

**Author notes:** Fernandes and Tellez-Colmenares contributed equally to this paper.

## Abstract

A substantial gap in dominance hierarchies research is understanding how such relationships change over time. Previous studies have generally used a single interaction matrix, assuming that dominance is temporally consistent. Yet food sources can be scarce and temporary, leading to changes in species frequency and assemblage composition. Hummingbird assemblages fluctuate as these tiny birds track seasonal floral resources, with individual movements affecting species composition. Artificial feeders, which may foster a greater number of aggressive interactions, provide a stable environment for studying dominance, unlike ephemeral and shifting wild resources. To assess the temporal stability of dominance relationships, we documented aggressive interactions in a hummingbird assemblage over three months on the western slope of the eastern Cordillera of Colombian Andes. We tracked local changes in species composition and visit frequency, recording interactions among 11 hummingbird species using high-speed cameras at three time scales: hour, day, and month. Dominance relationships and their certainty were analyzed through Perc analyses and interaction networks. Over three months of recording, and perusing over 27 hours of non continuous highspeed videos, we registered 3,907 aggressive interactions and 12,629 visits. Our results revealed substantial temporal variation in dominance relationships, species composition, and visit frequency, even within the constant food source that feeders provide. Dominance hierarchies were generally intransitive, with large species dominating, but also with smaller species occasionally winning interactions. We found high uncertainty in dominance relationships within shorter intervals with fewer interactions (*e.g*., hourly); and that rank changes between dominant species across months and days were linked to shifts in feeder visits. We recommend that future studies incorporate temporal variation, certainty analyses, and interaction networks to determine whether dominance relationships are stable enough to form reliable hierarchies and to explore the ecological significance of reversals in dominance and associated determinants.

**Highlights:**

- Dominance fluctuates across temporal scales, with monthly, daily, and hourly variations.
- Hierarchies become less stable at finer temporal scales.
- Body size and feeder visitation frequency shape dominance interactions.
- Smaller species initiate more aggressive interactions but lose more frequently.
- Relationships among dominant species are less predictable than among subordinates.

## 1. Introduction

Competition is a fundamental evolutionary force shaping ecological assemblages, behavioral ecology and population dynamics. To compete for resources, animals can either deplete them before their competitors (exploitation competition), or rely on aggressive behaviors to physically exclude contenders (interference competition) (Rico-Guevara et al., 2019 [1] ; Sargent et al., 2021[2] ; Valenzuela-Toro et al., 2023[3] ; Vard et al., 2002[4]). Animals that rely on exploitation strategies do not fight for food access but must be capable of quickly reaching and consuming the available resources (Valenzuela-Toro et al., 2023[3]). (Conversely interference strategies allow for monopolization (*e*.*g*., of a food patch) but can come at high costs given the time and energy spent on aggressive displacements and the elevated risk of injury that comes with physical confrontation (Brown, 1964[5]). Studying aggressive interactions among species that exploit the same resources is therefore pivotal to understanding competition dynamics. Identifying winners and losers in such interactions allows us to define dominance patterns within social groups or species assemblages. Multiple aggressive interactions among individuals establish dominance hierarchies (Altshuler et al., 2004[6] ; Carpenter, 1978[7] ; Gowda et al., 2012[8] ; Temeles et al., 2004[9]) which emerge through agonistic behaviors and differences in competitive abilities (Adams, 2001[10] ; Smith and Parker, 1976[11]). Although aggression entails energetic costs and potential risks of injury, individuals that win such contests typically gain priority access to resources, mates, or territories–benefits that can outweigh those costs. Moreover, repeated victories can build a dominance reputation, reducing the need for future fights and increasing overall fitness payoffs Tibbetts et al., 2022[12]). Yet, how stable these dominance hierarchies remain through time is less understood. Do individuals or species retain their dominant status once established, or must they continually reassert it as environmental and social conditions change? The permanence–or instability– of dominance relationships has important implications for the evolutionary pressures shaping competitive behavior and the maintenance of social order in animal communities (Tibbetts et al. 2022[12]).

The temporal dynamics of these hierarchies, however–whether they remain fixed or shift over time–remain underexplored. Behavioral research has demonstrated that dominance hierarchies are not fixed, but can change due to factors such as age, experience, social interactions, and environmental fluctuations (Bedford et al. 2017[13] ; Francis et al., 2018[14] ; Márquez-Luna et al., 2019[15]). For example, in rhesus macaques (*Macaca mulatta*) dominance relationships can change as individuals age and as the social structure of the group evolves, with older and more experienced individuals often occupying higher ranks (Tiebout, 1996[16]). Similarly, in wolves (*Canis lupus*), pack hierarchies fluctuate depending on physical condition and social alliances of the individuals involved (Mech, 1999[17]). In social insects like ants, interspecific dominance hierarchies vary based on species’ relative abundance, the outcomes of aggressive interactions, and their combined effects (Davidson, 1998[18 ; Stuble et al., 2017[19]). These examples illustrate that dominance is a dynamic property shaped by both internal and external factors. However, despite extensive work across taxa, the temporal persistence of these hierarchies–the extent to which dominance relationships remain stable or reorganize through time–remains poorly understood. Understanding such temporal shifts has powerful implications for interpreting social structure and the ecological consequences of competition.

Hummingbirds provide an ideal system for studying changing dominance hierarchies. These birds exhibit frequent, intense aggressive interactions (displays, chases, and direct attacks; Camfield, 2006[20] ; Rico-Guevara and Araya-Salas, 2015[21]) often influenced by factors like body size (Echevarría and Buitrón-Jurado 2025[22] ; Altshuler et al., 2004[6] ; Justino et al., 2012[23 ; Lara et al., 2009[24] ; Stiles and Wolf, 1970[25]), physical status (González-Gómez and Vásquez 2006[26] ; Kodric-Brown and Brown, 1978[27]) age Rousseu et al., 2014[28]), wing morphology (Feinsinger and Chaplin, 1975[29] ; Feinsinger and Colwell 1978[30]) and flight aerodynamics (Altshuler, 2006[31]). Larger hummingbird species tend to displace smaller ones from high-quality resources, whereas smaller species often compensate by feeding at different times of day, in peripheral patches, or on lower-quality resources to avoid competition (Justino et al., 2012[23] ; Lara et al., 2009[24] ; Stiles and Wolf, 1970[25]). The intensity of territoriality and competition is also regulated by the number of competitors (Stiles and Wolf, 1970[25] ; Wolff, 1985[32]) and the quality of available resources (Rousseu et al., 2014[28]). Consequently, hummingbird dominance hierarchies can shift significantly not only in response to resource availability (Grant, 1994[33] ; Wolf, 1978[34]), individual traits (*e*.*g*., size age), relative abundance of competitors who differentially track blooming of profitable plant species which they can efficiently exploit (Gutiérrez et al., 2004[35] ; Rico-Guevara 2021[36] ; Rodríguez-Flores and Stiles, 2021[37]) and other social interactions.

Despite this known variability in hummingbird dominance hierarchies, their temporal dynamics across fine and broad time scales has received little attention (Letourneur, 2000[38] ; Troen et al., 2008[39]). While dominance is closely linked to access to food and nesting sites, as well as changes in species abundance and assemblages composition that often correlate with resource availability (Lyon, 1973[40] ; Maher and Lott, 2000[41] ; Stiles and Wolf, 1970[25] ; Stuble et al., 2017[19]) dominance is not determined solely by these coarse-scale processes. Fine-scale temporal variation–hourly, daily, or sub-seasonal–may reflect mechanisms that remain invisible when examined only through seasonal breeding or flowering cycles. For example, short-term changes in energy balance, competitive motivation, or transient dominance reversals may influence which individuals or species gain access to feeders or flowers during specific periods of peak activity (Landau, 1975[42]). Yet these short-term dynamics remain largely undocumented in natural hummingbird assemblages (Miller et al., 2017[43] ; Fernandes et al., in prep).

Previous research has shown that dominance can change with nectar supply Kodric-Brown and Brown, 1978[27]), but these studies were restricted to partially controlled conditions or focused on coarse comparisons across flower densities. What is still missing is a clear understanding of how dominance fluctuates at the temporal scales at which hummingbirds actually make foraging decisions–minutes to hours–and how such short-term shifts accumulate into longer-term patterns (Bukacińska and Bukaciński 1994[44]). Aggression may vary with energetic state, time of day, individual turnover, transient resource pulses, or asymmetric motivation unrelated to breeding cycles (*e*.*g*., territorial replacements, juvenile recruitment), yet no empirical framework has evaluated these dynamics in the wild.

In tropical systems–where many hummingbirds live–breeding cycles are weakly seasonal or occur year-round. Therefore, dominance hierarchies may not be anchored to predictable circannual rhythms, making shorter-term fluctuations especially important. Understanding temporal variation therefore becomes especially relevant: without clearly defined breeding seasons, dominance may be shaped by shorter-term fluctuations in resource pulses, rainfall events or social turnover processes that remain unexplored. Thus, examining temporal variation at multiple scales allows us to disentangle dominance shifts driven by breeding or seasonal flowering peaks from those emerging spontaneously from the competitive system itself.

To address these gaps, artificial feeders provide a powerful system in which the temporal dynamics of dominance can be examined under controlled yet ecologically relevant conditions. Feeders concentrate multiple individuals and species in a small area, generating interaction rates similar to those observed at naturally occurring high-value nectar patches. In the wild, large flowering events of generalist plants–particularly mass-blooming trees such as many legumes–also attract dense, mixed-species assemblages of hummingbirds, where frequent chases, displacements, and aggressive encounters (Camfield, 2006[20]; Rousseu et al., 2014[28])unfold much like those documented at feeders. Thus, although artificial, feeders effectively recreate the social and competitive environment of naturally super-productive floral patches, making them a valuable context for studying dominance dynamics that are difficult to observe at dispersed or ephemeral flowers (Camfield, 2006[20]; Rousseu et al., 2014[28]). Moreover, as supplemental feeding becomes increasingly common across human-modified landscapes, understanding how interspecific interactions at feeders shape dominance hierarchies and aggressive behavior has growing ecological relevance in its own right (Avalos et al., 2012[45]). Artificial feeders, therefore, offer a tractable and ecologically realistic setting to quantify dominance interactions at fine temporal scales.

In our study, we explored the dynamics of interspecific aggressive interactions within a hummingbird assemblage using artificial feeders at three temporal scales: hourly, daily, and monthly. By conducting observations over three hours, for three days, and during three months, we quantified short-term shifts in dominance ranks and species composition. We analyzed these relationships through ranking indices and interaction networks.

We hypothesized that larger hummingbird species would generally domi-nate smaller ones because body size strongly predicts competitive advantage in hummingbirds through greater energy reserves, higher fighting ability, and more effective territorial defense. However, we also expected dominance to be non-linear (intransitive) rather than fixed-that is, smaller species could occasionally win some confrontations. This expectation is supported by previous evidence showing that dominance outcomes can depend on factors other than size, such as fluctuating nectar availability, individual motivation, energetic state, prior residency, or behavioral strategies that vary over time. Such context-dependent mechanisms can produce temporary reversals in competitive outcomes.

Additionally, we predicted that dominance relationships would vary over time (within months), similar to changes in species composition and feeder visitation. At the finest temporal scales, where community structure remains relatively stable (*e*.*g*., within hours), we expected dominance relationships to remain mostly consistent. By understanding how dominance hierarchies fluctuate, we can gain insight into the behavioral strategies and social structures that underpin hummingbird populations, with broader implications for the study of competition and dominance across ecological assemblages.

## 2. Methods

### 2.1. Study Site

This project was conducted at the Colibrí Gorriazul Research Station on the western slope of the Eastern Cordillera in Cundinamarca, Colombia (04^°^ 23’ N, 74^°^ 21’ W; 1,712 m.a.s.l.). This site includes a large diversity of hummingbird-pollinated plants and trees, fostering an ideal environment for studying hummingbird behavior. The station has maintained hummingbird feeders since 2013, with 15 feeders present filled with 18-20% Brix sucrose solution) when this project was developed in 2016. The feeders were strategically placed throughout an area of around 300 m^2^ to maximize observation opportunities (see Ethical Note below for more details about the feeder setup).

These feeders attract an assemblage of 11 common species of hummingbirds (Table 1). he station’s consistent and controlled provision of resources creates a unique opportunity to observe and analyze the dynamics of dominance hierarchies and interspecific interactions among these hummingbird species. Additionally, the station’s location within a diverse ecosystem, rich in floral resources, allows for the examination of resource competition in a semi-natural setting.

**Table 1:** English and Latin names of the species used in this study, along with their corresponding species codes (which will be used for figures and tables throughout the paper). The average masses used in our analysis were taken from Hilty (2003)[121], with the exception of *Saucerottia cyanifrons*, for which we used Ayerbe-Quiñones (2024)[122].

| English name | Latin name | Species code | Mass (g) |
| --- | --- | --- | --- |
| Sparkling Violetear | *Colibri coruscans* | CC | 8.5 |
| White-necked Jacobin | *Florisuga mellivora* | FM | 8.0 |
| White-vented Plumeleteer | *Chalybura buffonii* | CB | 7.3 |
| Black-throated Mango | *Anthracothorax nigricollis* | AN | 7.0 |
| Brown Violetear | *Colibri delphinae* | CD | 7.0 |
| Green Hermit | *Phaethornis guy* | PG | 6.0 |
| Andean Emerald | *Uranomitra franciae* | UF | 5.3 |
| Lesser Violetear | *Colibri cyanotus* | CCY | 5.0 |
| Rufous-tailed Hummingbird | *Amazilia tzacatl* | AT | 5.0 |
| Indigo-capped Hummingbird | *Saucerottia cyanifrons* | SC | 5.0 |
| White-bellied Woodstar | *Chaetocercus mulsant* | CM | 4.0 |

### 2.2. Aggressive Interactions

We identified hummingbirds to species and age group (we include only adults) but not by sex, as most species were monochromatic or indistinguishable in the recordings. We quantify interactions and feeder visits using JVC GC-PX100 high-speed camcorders (240 frames per second), positioned approximately one meter away from the focal feeder to maintain a consistent field of view and to reduce disturbance (continuously recording for the entire trial period without a human operator present). High-speed recordings were conducted for 20-minute intervals starting at 800 hours, 1200 hours, and 1700 hours, on three days each month, across three months (March, May, and July, 2016), producing a total dataset of 27 hours of recordings. These time intervals were chosen to represent different levels of hummingbird activity throughout the day (based on preliminary data collection, Tellez-Colmenares 2018[46]). Each day, we recorded at three randomly selected feeders out of the 15 available in the study area. The resulting videos were analyzed frame-by-frame using QuickTime Player 7.7.6 to identify interactions. These months differ in weather conditions (Berry and Steyermark, 1985[47]): rainy in March and May, while dry in July (Ecológicó-Ecoforest 2006[48] ; Tellez-Colmenares, 2018[46]).

We focused on agonistic interactions such as chases and direct aggressions. For each observed aggression, we recorded the aggressor species (initiator) and the recipient species (target of the aggression). We recorded the outcome of each interaction from the aggressor’s perspective rather than the recipient’s (*e*.*g*., if A attacked B, the outcome was categorized with respect to A), to maintain consistency throughout our analyses. Interaction outcomes were then classified as one of three mutually exclusive categories: victories (when an individual successfully distances its opponent from the resource or prevents other individual from approaching the resource), losses (when an individual is unable to access the resource due to being driven away or prevented from reaching it), and ties (when both individuals feed simultaneously post-interaction); this resulted in a matrix of discrete data from individual-to-individual interactions, but tabulated at the species level. Intraspecific interactions were included solely for descriptive analyses, because we only identified winners/losers at the species level. Additionally, we recorded the number of approaching visits to each feeder by species (arrivals within the frame, regardless of whether they fed or not) throughout the entire 20-minute interval.

### 2.3. Dominance Analysis

The descriptive analysis involved a comprehensive examination of various parameters, including total initiated interactions (started aggressions), interspecific victories and losses, intraspecific aggressions, and recorded visits. For further descriptive analysis per species in each month and day (Table S1), we calculated the ratio of losses to initiated interactions (total losses / total started aggressions) and the ratio of initiated aggressions per visit (total started aggressions / total visits).

All recorded behaviors, excluding ties (because they did not provide information of hierarchy level), were organized into matrices wherein “wins” were assigned to rows, “losses” to columns, and the diagonals represented intraspecific interactions (*i*.*e*. adjacency matrix). These data characterized the network structure, wherein species (nodes) interacted with other species through directed links. We built separate matrices for each month, day, and hour and compared network structures to explore potential temporal dynamics.

We used a network-based approach to calculate species’ dominance hierarchy, ranking order and dominance certainty, incorporating multiple indirect dominance pathways to quantify dyadic dominance relationships. This analysis was performed using the *Perc* package in R (Fushing et al., 2011[49] ; R Core Team, 2023[50]). The Perc index evaluates dominance through network percolation, a process that traces how dominance relationships spread through both direct and indirect interactions in a social network. In other words, it assesses not only who dominates whom directly but also how these effects propagate through others in the group. By integrating these pathways, the index provides a comprehensive view of dominance hierarchies, reflecting the overall flow of dominance within the network. The importance of this approach lies in its ability to reflect the complexity of social interactions by considering both direct and indirect relationships, providing a more accurate representation of dominance hierarchies. We only included species with dominance certainty ≥ 0.85 which ensures that the inferred dominance relationships are robust and not driven by sparse or ambiguous interactions.

All network measures and visualizations were conducted in R using the *igraph* v0.7.1 (Csárdi et al., 2025[51]) and *Sna* v2.3-2 Butts 2008[52]) packages. We applied two standard measurements to compare network structural similarities and differences:

1. Density: This metric measures how well-connected the network is, indicating the proportion of observed interactions out of all potential interactions among species (Wasserman and Faust, 1994[53]). The density index ranges from 0 to 1, with 1 representing a fully connected network. A higher density suggests more frequent interactions and a potentially more cohesive social structure. We tested whether the density of each network statistically differed from a theoretically maximally connected network (density = 1) using bootstrapping methods provided by UCINET (So et al., 2015[54]). Ve compared densities between networks using only species present in all three months of the study (March, May, July). Within months, comparisons between days were restricted to species observed consistently across all three sampled days, employing a bootstrapped version of the paired t-test with 5,000 permutations in UCINET. Density aids in understanding the intensity of social interactions which can impact resource allocation and species coexistence.
2. Reciprocity: This metric represents the proportion of mutual ties in the network (Wasserman and Faust, 1994[53]). Higher reciprocity indicates that species tend to reciprocate aggressive interactions, which could signify balanced power dynamics within the assemblage. We analyzed the variability of aggressive interactions across species in the network, testing whether the network had significantly higher reciprocity than expected by comparing it to 5,000 random networks with the same number of nodes and graph density. Understanding reciprocity is biologically important as it provides insights into the stability and mutuality of social relationships, which can influence group cohesion and cooperation.

To compare the dominance rank positions of common species among hours, days, and months, we used the following indices:

1. Out-degree and In-degree: These metrics represent the number of ties from (out-degree) and to (in-degree) each species (Hanneman and Riddle, 2005[55]). Out-degree measures how many other species with which a particular species initiates interactions, while in-degree measures how many species initiate interactions towards a particular species. High out-degree might indicate a dominant species frequently engaging in aggressive behaviors, whereas high in-degree could indicate a species that is frequently targeted. We tested whether the maximum out-degree of each network significantly differed from random networks with the same number of nodes and graph density using the *sna* package in R. We did not test in-degree for significance because be-ing the target of aggression can be strongly influenced by factors unrelated to dominance (*e*.*g*., abundance feeder position). Nevertheless we report in-degree values descriptively to complement out-degree and provide a fuller picture of species’ network positions.
2. Hub and Authority Centrality (Kleinberg 1999[56]): Hub centrality measures the “importance” of a species based on the number of outgoing links, weighted by the authority of the species they target. Authority centrality measures the importance of a species based on the number of incoming links, weighted by the hub scores of the aggressors. A species can have a high outdegree but low hub centrality if it mostly targets species with low authority, while another may have low in-degree but high authority if the aggressions it receives are predominantly from influential hubs. These centrality measures are thus complementary to degree metrics and crucial for understanding the roles different species play in the network which can provide information about their competitive strategies and social dominance.

We tested the significance of network density differences using bootstrapping methods provided by UCINET (Borgatti et al., 2002[57] ; So et al., 2015[54]). Ve compared densities among networks using only species present in all three months of the study (March, May, July). Within months, comparisons between days were restricted to species observed consistently across all three sampled days, employing a bootstrapped paired t-test with 5,000 permutations in UCINET (Borgatti et al., 2002[57]). The patterning of relationships between species across different months and days was compared using the quadratic assignment procedure (QAP) regression in UCINET (Krackhardt 1987[58] ; So et al., 2015[54]). QAP regression computes standard regression across all species cells of the behavior matrix, then randomly permutes the cells and repeats the regression 5,000 times. The observed regression coefficient is compared to the distribution of coefficients to determine a p-value (So et al., 2015[54]). The biological importance of these tests lies in identifying consistent patterns and temporal shifts in dominance hierarchies which can reflect changes in environmental conditions, resource availability, and social dynamics.

### 2.4. Ethical Note

This study adhered to ethical standards and legal requirements while providing valuable insights into hummingbird behavior in a semi-natural setting. The feeders were filled with an 18-20% Brix sucrose solution which is within the range of nectar concentrations available in many hummingbird-pollinated flowers (*e*.*g*., Baker 1975[59] ; Brown and Kodric-Brown, 1979[60]; Stiles, 1976 [61]). The sucrose concentration was measured using a handheld refractometer (ATAGO, Master-53T; Brix 0.0 to 53.0%). The feeders were regularly cleaned and refilled to ensure a consistent food supply and maintain hygienic conditions, minimizing the risk of disease transmission among the birds (Wilcoxen et al., 2015[62]). No hummingbirds were captured or held in captivity, and all interactions were recorded without interfering with natural behavior. No invasive or harmful techniques were used; all species identification was based on observation. All filming activities were reviewed and authorized by the Institutional Animal Care and Use Committee at the University of Connecticut; Exemption Number E09-010.

## 3. Results

We collected data for 11 species of hummingbirds, observing 3 907 aggressive interactions (3,647 excluding ties), of which 40.25% were intraspecific, and 12,629 feeder visits. The composition and interactions varied across time scales – hourly, daily, and monthly – reflecting the temporal dynamics of dominance relationships (Figure 1, Table S1). Across all months, *Saucerottia cyanifrons* and *Amazilia tzacatl* were the most abundant, and thus initiated the highest number of aggressive interactions overall (Table S1). *Saucerottia cyanifrons* and *Amazilia tzacatl* also initiated intraspecific interactions more often than other species (57.03-87.63%, Table S1). Conversely *Chaly-bura bu_onii, Anthracothorax nigricollis, Colibri cyanotus*, initiated fewer than 20% intraspecific interactions on both daily and monthly scales. Regarding interspecific aggressive interactions, *S. cyanifrons, A. tzacatl*, and occasionally *C. cyanotus* experienced a higher amount of losses when initiating interactions (Table S1). *A. nigricollis* and *C. buffonii* displayed a greater proportion of interspecific aggressive interactions overall while *S. cyanifrons* exhibited fewer aggressions from heterospecifics per visit. Uncommon species such as *Uranomitra franciae* and *Phaethornis guy* rarely initiated interactions and *F. mellivora* and *Chaetocercus mulsant* were mostly absent.

**Figure 1.**
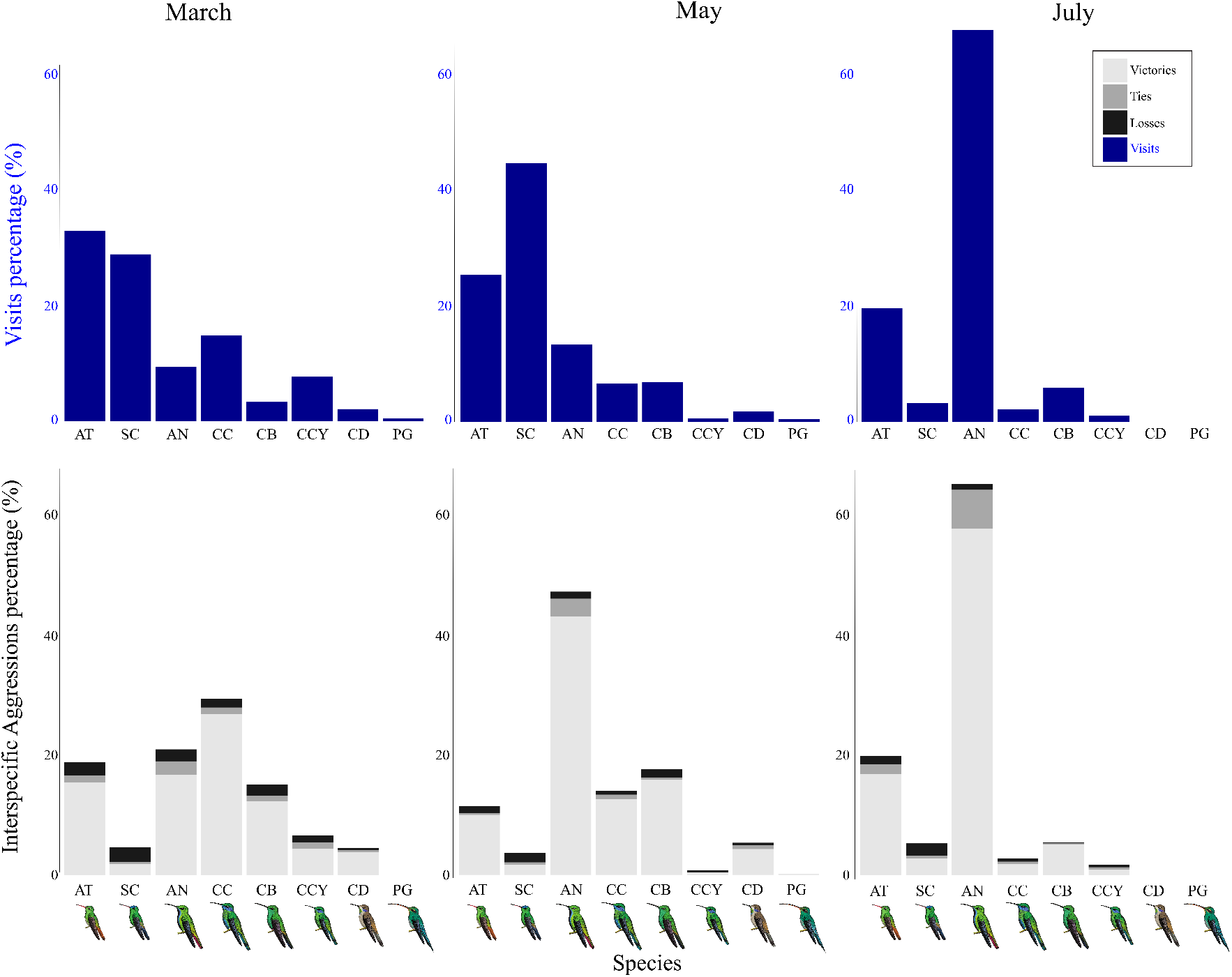
Monthly variation in the percentage of feeder-approaching visits (upper panel; see Table 1 for species codes) and interspecific aggression (victories, ties, and losses; lower panel) by the most common hummingbird species. Species are organized by approaching visits frequency, with this order maintained across all panels. Seven species visit the feeder most frequently throughout the month (with variation in the number of visits between months), while the others are considered rare visitors. The sizes of the hummingbird drawings are scaled relative to the bird’s actual size. Generally, species with more feeder-approaching visits tend to have more interactions (see *A. nigri-collis* AN; Table S1). Species’ drawings provided by Fernando Ayerbe-Quiñ ones.

The number of interspecific ties was generally low for all species except for *A. nigricollis* and *A. tzacatl*. The species with the most frequent tying was *A. nigricollis*, which consistently had a high number of ties at both the daily and monthly scales (Figure 1; Table S1). *A. tzacatl* also had a high number of ties, except in May. In contrast, despite being a frequent visitor, *S. cyanifrons* rarely had interspecific ties. In addition, March accumulated 49.6% of all recorded ties (most of them occurring on March 6), double of the other months. Regarding hourly distribution, 50.4% of ties occurred at 1700 hours, while the remaining ties were evenly distributed between 800 and 1200 hours, each registering approximately 25% Table S1).

The interaction data were compiled into a single interaction matrix, which was characterized by high density (0.875) and reciprocity (0.897) indicating a well-connected network with bidirectional interactions, but with low out-degree centralization, suggesting that dominance was not highly skewed toward a few individuals. Most species exhibited similar out-degree values (Table S2), in-degree, in-closeness (Table S3) and out-closeness values (Table S4) except for *P. guy*, which had the lowest out-degree and hub scores, indicating limited success in aggressive interactions. In contrast, *Colibri delphinae* had the lowest betweenness centrality (Tables S5-S6), while *S. cyanifrons* recorded the highest, highlighting its pivotal role in connecting interaction networks.

Dominance hierarchy based on the total interaction matrix reflected that larger and more aggressive species, such as *C. coruscans* and *A. nigricollis*, occupied the highest ranks in the dominance hierarchy, while smaller but more frequent species like *A. tzacatl* and *S. cyanifrons*, ranked lower. *P*.*guy*, a rare and non-territorial species ranked at the bottom of the hierarchy (Figure S1).

### 3.1. Dominance Across Months

Temporal variation in dominance dynamics and interaction networks was evident at daily and hourly scales, and even more pronounced at the monthly scale (Figures 2, S1-S3). Dominant species such as *C. delphinae, C. buffonii*, and *Colibri coruscans* exhibited seasonal shifts, with higher aggression in March and May but lower in July. In contrast, subordinate species like *S. cyanifrons* and *A. tzacatl* maintained stable relative visit frequencies across months although absolute visitation decreased markedly for all species in July (Figures 2-3 and S1).

**Figure 2.**
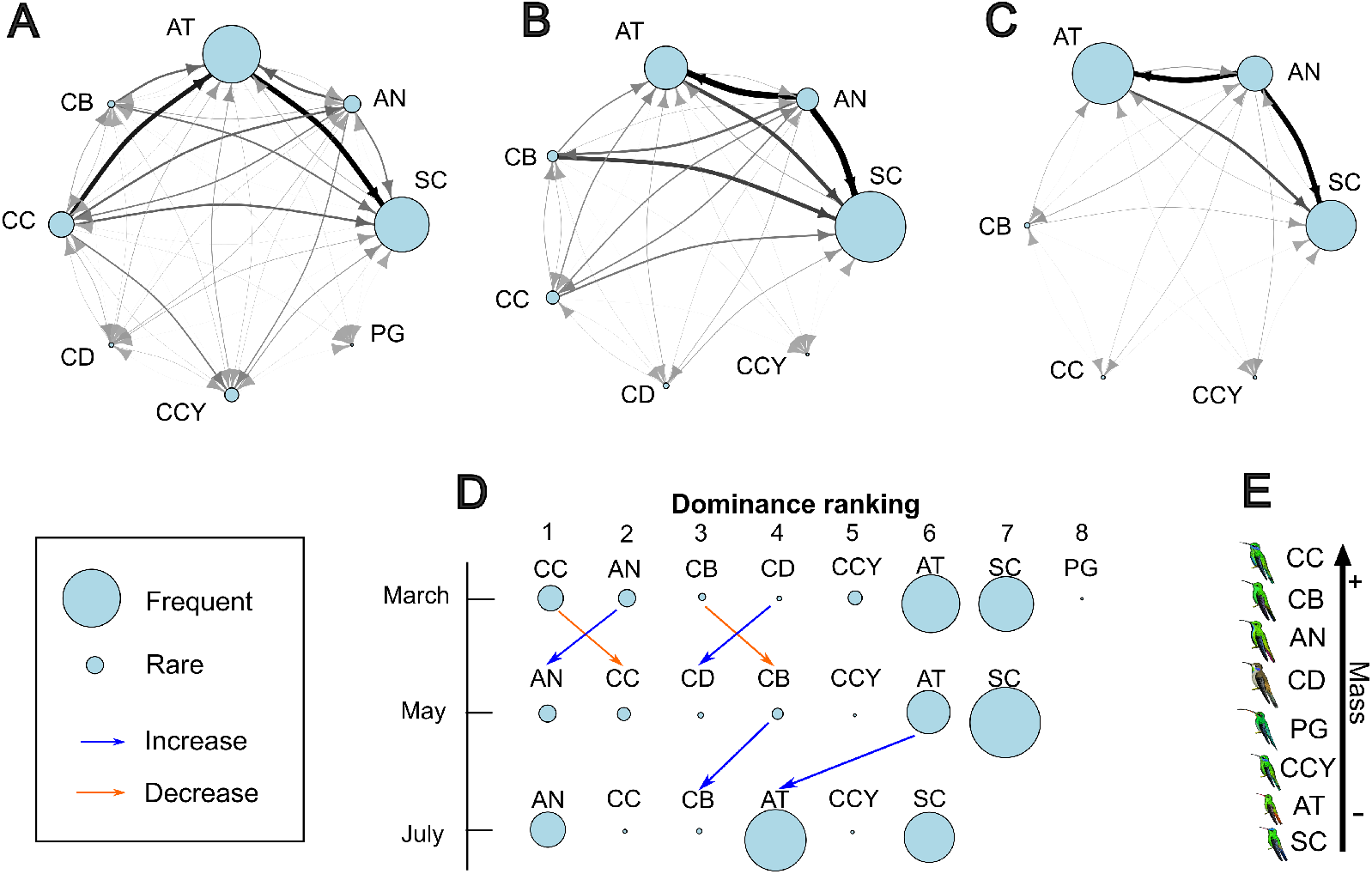
Dominance variation across months. Node size in the interaction network corresponds to species frequency (relative visits to the feeder; See Table S1). Darker and thicker links indicate a higher number of interactions between those species in the arrow direction. The dominance hierarchy is determined according to the Perc method, displaying only species with high certainty levels of dominance (dominance certainty ≥ 0.85). Blue arrows represent a ranking increase, whereas orange arrows indicate a rank decrease. A) March network; B) May network; C) July network; D) Dominance ranking variation across months; E) Species’ body mass scale. See Table 1 for the codes of hummingbird species. Note that node sizes for a given species–particularly the most common visitors, AT, AN, and SC–can differ across months. Also note the changes in dominance ranking of the species among months and how it correlates with frequency of visits (*e*.*g*., CC and AN in March and May). However, when the difference in mass is not bigger (Table 1), the changes in ranking sometimes can occur if the visit frequency difference is very high (see CCY and AT in May and July). Species’ drawings provided by Fernando Ayerbe-Quiñones.

**Figure 3.**
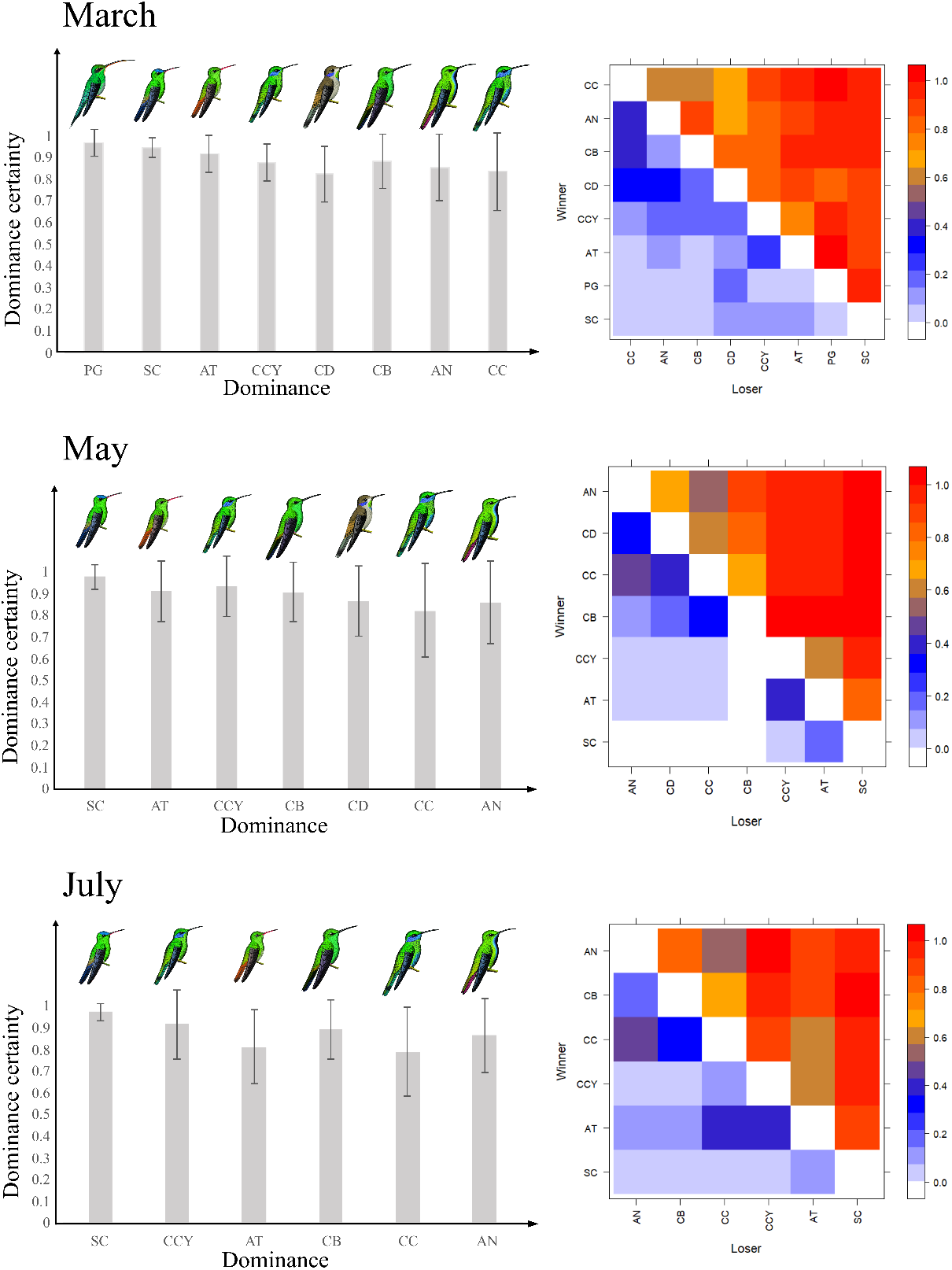
Dominance certainty across months. Dominance certainty was higher in the species with lower ranks; note the value of dominance certainty on the left side of the image and the dominance probability heatmap on the right, which depicts the simulated probability of being ranked subordinate or dominant based on our data (with the X axis showing losing species, and the Y axis showing winning species). The heatmap’s color scale signifies clearly defined relationships in which a species is linearly ranked as subordinate (blue region) or dominant (orange-red region), with the exception of when there were no data (white, 0.0, signifying intraspecific interactions), or when the dominance rank was unclear (brown region, ∼0.5).

Dominance shifts appeared linked to visit frequency for some species. For example *C. coruscans* ranked high when they also exhibited higher visitation but declined with reduced visits, whereas *A. nigricollis* could surpass *C*.*coruscans* in the dominance rank when the former species is more frequent. However *C. buffonii, C. delphinae*, and *C. cyanotus* showed no clear link between visit frequency and dominance. Subordinate species such as *A. tzacat* and *S. cyanifrons* consistently maintained lower dominance ranks regardless of their visit frequency. At finer temporal scales, no consistent hourly or daily dominance patterns emerged.

March exhibited the highest network density and reciprocity, suggesting that interactions were frequent and often bidirectional. In contrast, May showed the greatest out-degree and path length, indicating that dominance interactions were more hierarchical but not strongly polarized; several species occupied mid-rank positions rather than clustering at the extremes of the hierarchy (Tables S7–S11). In March, most species had equal hub and authority values (Tables S7–S8) meaning they experienced a similar number of wins and losses (intermediate dominance). Species such as *S. cyanifrons* had the highest in-closeness in March, suggesting frequent interactions, likely driven by its high visit frequency (Figure 3). By May, *S. cyanifrons, C. cyanotus*, and *A. tzacatl* showed the lowest out-degree and out-closeness, driven by having fewer winning interactions. Meanwhile, *C. buffonii* and *C. delphinae* had the lowest in-degree and in-closeness implying they were rarely targeted in agonistic heterospecific interactions. Additionally, *S. cyanifrons* had the lowest hub value, indicating a higher frequency of losses, further reinforcing its lower dominance ranking. In July, *C. coruscans* initiated fewer interactions and had the lowest authority score, indicating that it acted as a weak competitor in these encounters and seldom defended or won interactions. *A. nigricollis* and *C. buffonii* (also dominant species) ranked highest in hub values indicating they won more interactions. Meanwhile, *S. cyanifrons* consistently had the highest authority score, suggesting that despite losing frequently, it remained highly engaged in the network, possibly due to its persistence in foraging attempts (high frequency of feeder visits) despite aggression from dominant species.

Across months, dominance certainty generally declined at finer temporal scales, although subordinate species exhibited stable high certainty values (Figures 3, S13-S16), indicating that their lower status was consistently reinforced over time. Monthly dominance indices generally lacked correlation, meaning that dominance relationships were not rigid and could fluctuate between months. However, out-closeness (March–July) and hubs (May–July) remained correlated, suggesting that certain species maintained consistent interaction patterns over these periods. Notably, dominance relationships were most stable between May and July (QAP; Table S11), suggesting that long-term dominance structures persisted despite short-term variations (Fig-ures S17–S25) and shifts of relative abundance among species Figure 1).

### 3.2. Dominance Across Days

March exhibited the highest day-to-day variation in dominance indices, particularly in network density and reciprocity (Table S5). On March 6, high density and reciprocity indicated frequent and bidirectional interactions, but low out-degree suggested that no single species dominated the network. In contrast on March 8, *C. coruscans* displayed the highest out-degree, indicating that it initiated most agonistic interactions that day. This reflects a more centralized dominance structure, where a single species temporarily acted as the main aggressor, in contrast with the more egalitarian interaction pattern observed on March 6. Over three days in March, dominance relationships correlated significantly (Table S12), with large species (*C. coruscans, A. nigricollis*) alternating as dominant, reflecting a dynamic but structured hierarchy. Meanwhile, S. cyanifrons consistently occupied the lowest rank despite being a frequent visitor at the feeders (Figure 4). Because each daily dataset was based on three 20-min observation periods, the absence of a species on a given day may reflect short-term sampling rather than a true absence from the assemblage. Unlike the patterns observed across months – where shifts in dominance were closely associated with species abundance– day-to-day changes appeared decoupled from abundance, suggesting that daily dominance dynamics were more stochastic and influenced by short-term behavioral fluctuations. Dominance certainty varied by day, with lower-ranked species showing greater stability on March 6 while dominant species exhibited more consistent values on March 9 (Figure S14). This contrasts with the monthly networks, in which dominance certainty was consistently high for all species.

**Figure 4.**
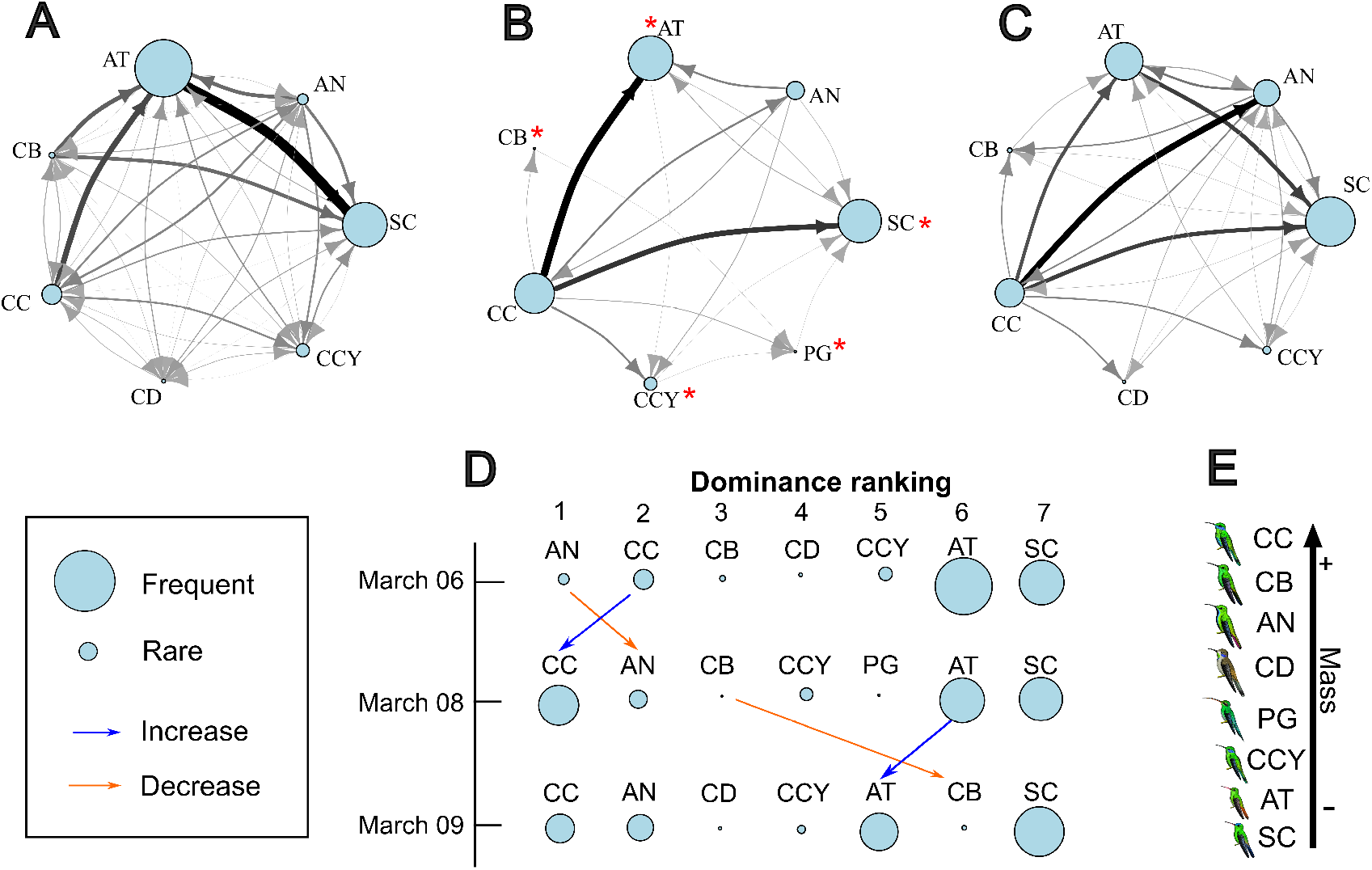
Dominance variation across days, an example for March. Node size in the interaction network corresponds to species frequency (relative visits to the feeder; See Table S1). Darker and thicker links indicate a higher number of interactions between those species in the arrow direction. The dominance hierarchy is determined according to the Perc method, displaying only species with high certainty levels of dominance (dominance certainty ≥ 0.85). Blue arrows represent a ranking increase, whereas orange arrows indicate a rank decrease. A) March 06 network; B) March 08 network; C) March 09 network; D) Dominance ranking variation across days; E) Species’ body mass scale. See Table 1 for the codes of hummingbird species. Species marked with an asterisk (*) have very low dominance certainty, a pattern that contrasts with the much higher certainty observed in the monthly networks (Figure 2), where no species exhibited low-confidence rankings. Notably, day-to-day changes in dominance do not closely track species abundance, unlike the clearer abundance-dominance patterns observed across months; this suggests that dominance dynamics at the daily scale may be more stochastic or influenced by shortterm behavioral fluctuations. Note the changes in visit frequency, species composition and rankings, and changes in network connectivity (number of links between nodes) between days.

May was the most stable month, with strong correlations in dominance relationships across days (Table S13). *S. cyanifrons* consistently had the lowest out-degree but the highest in-degree and in-closeness, indicating that while it engaged in many interactions, it rarely won the aggressions (Figure S2). *A. nigricollis* and *C. coruscans* remained dominant, though their rankings fluctuated slightly across days. Less dominant, species (*A. tzacatl, S. cyanifrons*) consistently had higher dominance certainty (Figure S15). Stability in May was reflected in strong correlations between May 13-15 despite minor changes in species presence, such as the absence of *C. delphinae* on May 15 (Figure S15).

July displayed the most dynamic ranking shifts between days (Figure S3). *A. nigricollis* maintained a dominant position, whereas *C. coruscans* showed high variability, including an absence on July 10. Betweenness and hub values varied significantly, with *A. nigricollis* consistently ranking highest, reinforcing its strong influence within the dominance network. However, dominance certainty was highly variable and generally low, reflecting instability in hierarchical positioning.

Despite daily fluctuations, strong correlations in in-closeness and authority values between consecutive days in both May and July suggest that species interaction patterns remained relatively stable over short timescales. This persistence indicated that while dominance hierarchies fluctuated, species maintained consistent roles within the network, reinforcing a structured but flexible dominance system.

### 3.3. Hourly Variability in Aggressive Interactions

Dominance indices– including density, reciprocity out-degree, and indegree –varied throughout the day highlighting dynamic shifts in the structure of dominance relationships. However, these changes had a consistent daily pattern. Some species exhibited significant shifts in rank over the course of the day, while others maintained more stable positions. Notably additional species were observed in the late afternoon, especially around 1700 hours (Figure S4). On March 6, high density and reciprocity were observed at 800 hours, indicating a high frequency of mutual interactions. For instance *S. cyanifrons* consistently had low out-degree and high in-degree values, reflecting its subordinate but central role in the interaction network–it was often involved in conflicts but rarely won them and had limited influence over other species. In contrast, *C. buffonii* had the lowest in-degree and incloseness, suggesting it was rarely targeted and occupied a peripheral role in the network. Despite this peripheral role, *C. buffonii* showed greater stability in dominance rank early in the day (at 800 hours) while *S. cyanifrons* maintained relatively stable ranks across the day as a whole (Figure S16). In the late afternoon, networks included additional species, reshaping interaction patterns (Figure S4–S6). On March 8, *C. coruscans* maintained a stable dominant position, while *S. cyanifrons* and *A. tzacat* consistently occupied lower ranks. In contrast, March 9 exhibited greater instability, with significant variation in dominance ranks throughout the day. For example *S. cyanifrons* shifted between dominant and subordinate roles, possibly reflecting momentary advantages (*e*.*g*., fewer competitors present) or increased persistence in foraging attempts (Figure S6). However, many species did not engage in many interactions to reliably estimate their dominance (low dominance certainty; Figures S17–S19) indicating incomplete dominance relationship information.

Hourly analyses during May showed that dominance certainty was generally stable with less dominant species showing higher certainty values (Figure S15). Across both in May (Figures S7–S9 S20–22S) and July (Figures S10-S12 S23-S25), dominance rankings at the top of the hierarchy fluctuated considerably across hours, while mid-ranked species exhibited relatively stable positions. Although dominance indices varied significantly across hours some species maintained consistent interaction patterns. While dominance networks remained significantly correlated across dates (Table S12) the hourly fluctuations in rankings highlight the importance of fine-scale temporal resolution to capture the fluid nature of dominance interactions. These patterns suggest that hummingbird dominance is both structured and flexible shaped probably by immediate social and ecological contexts. Thus beyond assessing whether dominance rankings are stable or not, examining variation at this fine temporal scale allows us to detect subtle, yet consistent within-day shifts in behavior that reshape the interaction network. In this sense hourly patterns reveal emergent behavioral structures that would remain hidden if dominance were analyzed only at broader temporal scales.

## 4. Discussion

### 4.1. Aggression Frequency

When food availability is extremely high, as observed at the Colibrí Gorriazul Research Station, aggression is also frequent (as seen in Carpenter and MacMillen 1976[63]; Goss-Custard 1970[64] ; Goss-Custard et al., 1984 65 ; Myers 1981[66] ; Myers et al., 1979[67]) consistent with the idea that high resource availability can sustain elevated levels of competition (as suggested by Maher and Lott 2000[41] ; Missagia and Castro-Verçoza 2014[68]). The high number of recorded aggressions surpasses those found in other studies conducted at both flowers and feeders (*e*.*g*., Armstrong 1991[69] ; Feinsinger 1976[70] ; Lee and Sung 2023[71] ; Lyon 1976[72] ; Lyon et al., 1977[73] ; Rousseu et al., 2014[28]) possibly due to the high number of feeders (15) and the prolonged availability of predictable food at the site.

Although species with more feeder visits interacted more frequently simply due to higher encounter rates, this did not translate into greater dom-inance. Several abundant species, such as *S. cyanifrons* engaged in many interactions but rarely won them, whereas less frequent visitors sometimes held higher dominance positions. Thus, abundance alone did not appear to determine aggressive performance in this assemblage. The species that visited the feeders most frequently were the smaller ones (*A. tzacat* and *S. cyanifrons*). The larger and more dominant species also visited them regularly, but there seemed to be fewer individuals of these species, or they visited less often. This pattern is consistent with the findings of Miller et al., (2017)[43] who studied seed feeders that attract species from many bird families and found that dominant species (*e*.*g*., woodpeckers) were rare within the assemblage. In our case all interacting species belonged to the same family (Trochilidae) which may have influenced the frequency and intensity of aggressive behaviors. Hummingbirds are all nectarivores and there is usually elevated competition over limited nectar resources when several species can access them. However, feeders are virtually unlimited resources and, on many occasions, hummingbirds visited without displaying aggression (although we did not quantify the frequency of this tolerance), while in other cases, aggressive interactions resulted in ties, where at the end of the confrontation both individuals accessed the feeder simultaneously. Temporal shifts in behavioral patterns as found in other hummingbird species by Lara et al., 2009[24]) and the occurrence of ties may contribute to the coexistence of multiple species.

4.2. Factors Affecting Hierarchy

There is a high interconnection in the networks, with almost all species interacting with each other (*e*.*g*., Miller et al., 2017[43]). When food sources are concentrated competition can be very intense at both intraspecific Feinsinger 1976[70] ; Márquez-Luna et al., 2015[74] ; Mendonça and Anjos 2006[75]) and interspecific Stiles and Wolf, 1970[25] ; Tiebout 1996[16]) levels as we found in our assemblage. Aggressive behavior in hummingbirds has been studied by quantifying the number and intensity of interactions and has been linked to the amount of competitors present (Araujo-Silva and Bessa, 2010[76] ; Justino et al., 2012[23] ; Lyon 1976[72] ; Lyon et al., 1977[73] ; Rousseu et al., 2014[28]) and the outcomes are partially explained by competitors’ characteristics. For instance, body mass is consistently one of the strongest predictors of dominance (Francis et al., 2018[14] ; Miller et al., 2017[43]). Other traits–including wing disc loading coloration and individual experience–also influence aggressive performance and resource holding potential (Carpenter et al., 1993[77]). Additionally physiological state can modulate dominance as sex steroids and body condition vary seasonally and affect territorial aggression levels González-Gómez et al., 2014[78]). Taken together these studies indicate that dominance arises from a combination of structural traits (*e*.*g*., size and wing morphology) and context-dependent physiological factors. Hierarchical relationships among dominant species are more variable than among subordinates; this aligns with findings in other hierarchical animal systems wherein dominant individuals face stronger competition from similarly ranked rivals (Lyon 1976[72] ; Araujo-Silva and Bessa 2010[76]). In contrast, subordinate species maintain stable lower positions and employ alternative competitive strategies (*e*.*g*., opportunistic feeding; Antunes 2003[79 ; Feinsinger and Colwell, 1978[30] ; Lara et al., 2009[24] ; Mendonça and Anjos 2006[75]).

In general, larger species tend to dominate smaller ones Camfield, 2006[20] ; Cotton 1998 80 ; Del Coro Arizmendi and Ornelas, 1990[81] ; Francis et al., 2018[14]; Funghi et al., 2015[82] ; Lara et al., 2009[24] ; Martin and Ghalambor 2014[83] ; Mendonça and Anjos, 2006[75] ; Stiles and Wolf 1970[25]). However in our study an exception occurred within the dominant group where this conventional pattern did not always hold. Smaller species also demonstrated dominance over larger ones. Ve found that *A. nigricollis* ≈ 7.0 g) despite being smaller than *C. coruscans* ≈ 8.5 g) sometimes held a higher dominance rank. These shifts occurred not only when *A. nigricollis* won direct confrontations against *C. coruscans* but also when it accumulated higher numbers of victories over other species present at a given temporal scale. Thus its relative position was influenced both by pairwise outcomes and by broader interaction patterns across the network. This anomaly in size-based dominance may be explained by a small difference in sizes between these two species and a possible correlation with visit frequency when analyzing networks at different temporal scales. At specific temporal scales *A. nigricollis* exhibited greater visit frequency than *C. coruscans* and other larger species, aligning with its elevated dominance status, as it has been shown in other hummingbird assemblages (Araujo-Silva and Bessa 2010[76]). Cases where smaller species dominate larger ones in direct interactions are uncommon but documented (*e*.*g*., Martin and Ghalambor 2014[83]). In contrast higher dominance rankings of smaller species relative to slightly larger ones can arise from higher encounter rates or greater persistence at resources (as reported by Araujo-Silva and Bessa, 2010[76]). Our results align more with this second pattern: the elevated ranking of *A. nigricollis* emerged primarily from its higher visit frequency and greater number of interactions rather than systematic dominance over *C. coruscans*.

The most subordinate species in our study *A. tzacat* ≈ (5.0 g) and *S. cyanifrons* ( ≈5.0 g), showed little variation in either the number of visits or dominance consistently ranked lower in the hierarchy and exhibited higher feeder visit rates than larger species (Figures 2-3). Additionally, other small species such as . *franciae* and *C. mulsant*, did not attack other species despite multiple visits (as registered in other small species by Bribiesca et al., 2019[84]). Some small hummingbirds act opportunistically, feeding on nectar sources undefended by dominant species (Feinsinger and Colwell 1978[30]), altering their resource access patterns and preferences (as evidenced in Tiebout 1996[16] ; Tellez-Colmenares and Rico-Guevara, 2023[85]). The lowest rank in the dominance hierarchy in the overall matrix was attributed to the trapliner species *P. guy* ≈ 6.0 g), which visited the feeders sparingly and displayed minimal aggression around food resources (Figure S1; seen in Colwell 1973[86] ; Feinsinger and Chaplin, 1975[29] ; Gill, 1988[87] ; Machado 2009[88]) preferring to visit high-quality non-defended flowers Feinsinger and Colwell 1978[30] ; Stiles and Wolf 1979[89]).

### 4.3. Temporal Variations

In our study species showed variation in agonistic activity, hierarchical ranking, and visitation patterns across all the temporal scales studied. The dominance hierarchy of the hummingbird assemblage at the study site was highly dynamic over certain temporal frames (as seen in other locations and species by Samuels and Gifford, 1997[90] ; Valderrábano-Ibarra et al., 2007[91] ; Villiamson et al., 2016[92]) in contrast to many studies suggesting that dominance hierarchies tend to remain stable Bedford et al., 2017[13] ; Cafazzo et al., 2016[93] ; So et al., 2015[54]). However we did not find consistent dominance relationships among species across the different temporal scales studied. It has been reported in domestic dogs that aggression decreases over time once the hierarchy is established (Van Kerkhove 2004[94]). However in this study, dominance patterns were highly dynamic and constantly changing (as seen by Williamson et al., 2016[92]). Although hierarchies could be established many individuals appeared willing to engage in agonistic interactions, and even very subordinate species/individuals may win several contests.

We observed that hierarchy and dominance ordinations in the hummingbird assemblage constantly varied across days within each month, with the most significant changes occurring among species with intermediate and high dominance ranks. Species at the lower ranks tended to remain in the same positions (*A. tzacat* and *S. cyanifrons*; similar to the results found by Bedford et al., 2017[13] ).

Daily temporal variations in dominance may stem from differences in competitive strategies or resource exploitation preferences (as seen in other studies *e*.*g*., Carpenter 1978[7] ; Carpenter et al., 1993[77] ; Dearborn 1998[95]) wherein certain species regulate access to resources at specific times of day. Additionally, variations in resource exploitation preferences and daily activity peaks may shape competitive interactions, influencing when and how species engage in aggressions. Temporal variations in dominance might be related to energetic constraints, as energy reserves influence aggressive behavior (Gesquiere et al., 2025[96]). Dominant species might defend resources more intensely in the morning when replenishing energy after fasting during the night, whereas subordinate species could maximize foraging at popular food sources later in the day (as demonstrated in other assemblages *e*.*g*., Antunes 2003[79]). These short-term dominance shifts may therefore reflect adaptive behavioral strategies.

The lack of consistent dominance relationships at finer temporal scales suggest that species modulate their behavior according to local conditions, energetic demands, or competitive pressure. For instance, hummingbirds are known to show two foraging activity peaks throughout the day – one in the morning and another in the late afternoon before roosting – to replenish energy reserves *e*.*g*., Lara et al., 2009[24] ; Shankar et al., 2019[97]). This tendency may help explain the high proportion of ties and aggressive interactions observed late in the day, as more individuals are active and competition for access to resources intensifies. These shifts could also be influenced by learning processes or varying levels of motivation to access and defend resources (Krebs 1982[98] ; Richards 2002[99] ; Stiles 1982[100] ; Tobias 1997[101] ; Vojczulanis-Jakubas and Araya-Salas 2023[102]).

At the smallest temporal scale, hourly aggression patterns matched those found by Biswas et al., (2014)[103] showing a progressive increase in aggressive interactions across the day. A high proportion of ties occurred during the 1700 hour block, when multiple individuals were feeding simultaneously at the feeders, likely to ensure access to the resource. However, we did not find a consistent pattern of variation among intermediate and high dominance ranks at the hourly scale.

When analyzing day-to-day variation within a month, changes in dominance were too stochastic and less pronounced than those observed among months. Nevertheless, some alternation between days suggests that day or hour-level sampling alone is insufficient to derive stable dominance relationships across species. However these finer temporal resolutions are essential to reveal how dominance fluctuates within days, uncovering short-term patterns that would be obscured in coarse temporal aggregations. As expected the highest number of interactions occurred among the most frequent feeder visitors. However, the hourly interaction network appeared to be well-connected (see density values) and with many reversals (see intransitivity and reciprocity values).

In fully natural settings, the dominance hierarchy can change during flowering seasons due to the temporal variation in the feeding roles in the assemblage (*e*.*g*., territorial) the local abundance of competitors (Pimm et al., 1985[104]) and the preference of hummingbird species to defend specific resources (Rico-Guevara et al., 2019[1] ; Sargent et al., 2021[2] ; Valenzuela-Toro et al., 2023[3] ; Vard et al., 2002[4]). Furthermore resource availability may influence the species composition of hummingbird assemblages (MalpicaPiieros et al., 2018[105]). In feeders, although the resource was constant, temporal variation in the hierarchy seemed to occur more rapidly, possibly reflecting the frequency of competitor visits to feeders.

### 4.4. Feeders Versus Natural Settings

Artificial feeders usage is a widespread practice and can influence hummingbird ecological dynamics (Echeverry-Galvis et al., 2024[106]) and evolutionary patterns (Alexandre et al., 2025[107]). Feeders provide an ideal setting for studying dominance (Miller et al., 2017[43]) because they allow for the collection of large amounts of data, which is useful for building robust interaction networks. In contrast, observations at flowers are limited by plant phenology visibility and the number of interacting individuals (Feinsinger 1976[70] ; Wolf et al., 1976[108]) as well as by the logistical challenges of setting up and operating camera equipment in heterogeneous and less-accessible natural environments.

Feeders are a predictable food source (Francis et al., 2018[14]; Tellez-Colmenares 2018[46]) which leads species to overlap in their foraging activity (Robb et al., 2008[109]). Additionally food at feeders can be perceived as higher quality, and individuals frequenting them may behave more aggressively than in the absence of feeders (Foltz et al., 2015[110] ; Niederhauser et al., 2021[111]). HHowever animals in these modified environments can exhibit behaviors that approximate those seen in natural matrices (Samuels and Gifford 1997[90]). In the studied assemblage, despite the high number of aggressions, the number of visits (particularly from those species with low hierarchical ranks) indicated simultaneous access to resources despite intense competition (as found in other assemblages by Lyon 1976[72] ; Stiles 1981[112] ; Wolf 1970[113] ).

### 4.5. Future Directions

The presence of individuals with different dominance abilities influences interaction outcomes and dominance hierarchies. However, due to the difficulty of marking hummingbirds, dominance dynamics have often been studied only at the species level, limiting insight into individual variation (Miller et al., 2017[43]). Ve recommend that future studies prioritize the evaluation of dominance hierarchies at the individual level. Incorporating individual identification–through marking or other methods–would allow for the inclusion of key demographic variables such as age and sex which can affect dominance status (*e*.*g*., Rousseu et al., 2014[28]) as well as morphological traits linked to resource holding potential and coloration that could signal these hierarchical ranks (as it has been done at the species level, *e*.*g*.,, Bribiesca et al., 2019[84] ; Fernandez-Duque et al., 2024[114] ; Márquez-Luna et al., 2019[15]). Additionally, individual-level tracking would enable more accurate estimations of abundance, as currently, there is no way to differentiate between a species/sex/age having more individuals vs. some individuals being around more. In this study, dominance was assessed in relation to visit frequency, but this parameter does not fully reflect the abundance of the species. Future studies should also consider a broader range of behavioral data including variation in the intensity of interactions (Copenhaver and Ewald 1980[115] ; Ewald and Carpenter 1978[116] ; Lyon 1976[72]; Lyon et al., 1977[73] ; Powers 1987[117]) as well as vocalizations and display behaviors (Camfield 2006[20]) which were not accounted for here due to methodological constraints but may also vary across individuals, environmental conditions, and over time. Extending recording windows would also provide deeper insight into temporal variation in dominance, capturing finescale shifts across hours, days, or seasons. When field time is limited, longer continuous recordings within a single day may reduce the likelihood of missing infrequent visitors, thereby improving the characterization of their rank and reducing the risk of false absences produced by short observation windows. Moreover, it is crucial to examine dominance across multiple seasons or throughout the year, as shifts in floral resource availability and species presence may lead to temporal changes in hummingbird dominance.

Finally, future studies could benefit from incorporating ecological variables such as nectar quality (*e*.*g*., Ewald and Orians, 1983[118] ; Maher and Lott, 2000[41]) spatial distribution (Adams, 2001[10] ; Maher and Lott, 2000 41]), food density, and habitat complexity (Eason and Stamps, 1992[119]). These factors, though not addressed in the present study, can shape the nature and outcome of dominance interactions (*e*.*g*., Ewald and Carpenter, 1978[116] ; Powers 1987[117] ; Stiles 1973[120]). By combining ecological, temporal, and individual-level perspectives, future work can build on our findings to clarify how dominance contributes to species coexistence, offering broader insights into the ecological dynamics of hummingbird communities.

## 5. Author Contributions

Nicolas Tellez-Colmenares: Conceptualization, Data collection, Formal analysis, Investigation, Data curation, Writing-original draft, Writing - review & editing, Visualization. Ana Melisa Fernandes: Conceptualization, Formal analysis, Methodology, Investigation, Data curation, Writing-original draft, Writing - review & editing, Visualization. F. Gary Stiles: Conceptualization, Investigation. Alejandro Rico-Guevara: Conceptualization, Funding acquisition, Writing - review & editing, Supervision.

## 6. Data Availability statement

Code for the dominance indices and interaction networks is available here: https://github.com/AnaMeliTucusito/Dominance-varies-at-different-time-scales-R-code-and-simulated-data-for-dominance-network-analysis. The data that support the findings of this study are available upon author (NTC) request.

## 8. Inclusion and Diversity statement

Our study includes scientists based in Colombia (NT, AMF, GS) where the study was conducted, and is composed of a mixed gender team who are residents of the local area in which the study was performed.

## 9. Acknowledgments

We are grateful to Jorge Pérez-Emá n for his insightful input on the initial idea for this manuscript and his invaluable support during the early stages of this work. We extend our sincere thanks to Fernando Ayerbe for generously sharing his beautiful illustrations. We also thank Mary, Lucero, and Parmenio– collaborators at the Colibrí Gorriazul Research Station– for their essential assistance during data collection. We appreciate the contributions of the Behavioral Ecophysics Lab at the University of Washington particularly Alyssa J. Sargent, as well as the anonymous reviewers, whose thoughtful feedback greatly enhanced the quality of this manuscript

