## supplements for "Temporal patterns of dominance in a hummingbird assemblage": Supplementary material.pdf

**Table S1. Aggression data across temporal scales (month, day, hour).** The table includes the following variables: intraspecific total started aggressions (number of intraspecific aggressions initiated by each species); % intraspecific aggressions (percentage of intraspecific aggressions relative to the total number of aggressions initiated); started interactions (total number of aggression events initiated by each species); losses if started (number of interactions lost when the species initiated the aggression); ties (number of interactions in which no clear winner was observed and both individuals accessed the resource); visits (number of arrivals to the feeders); proportion of losses vs. started aggressions (proportion of initiated interactions that resulted in a loss); and proportion of started aggressions per visit (proportion of initiated aggression events relative to the total number of visits by each species). * *Not started* indicates that a species did not initiate any interactions, making it impossible to calculate some indices.

**Conventions for Figures S1–S12.** Node size in the interaction network corresponds to species frequency (relative visits to the feeder; See Table S1). Darker and thicker links indicate a higher number of interactions between those species in the arrow direction. The dominance hierarchy is determined according to the Perc method, displaying only species with high certainty levels of dominance (dominance certainty ≥ 0.85). Blue arrows represent a ranking increase, whereas orange arrows indicate a rank decrease. Species with a * are those with very low certainty. Species codes (ordered from lightest to heaviest species): White-bellied Woodstar, *Chaetocercus mulsant* (CM); Indigo-capped Hummingbird, *Saucerottia cyanifrons* (SC); Rufous-tailed Hummingbird, *Amazilia tzacatl* (AT); Lesser Violetear, *Colibri cyanotus* (CCY); Andean Emerald, *Uranomitra franciae* (UF); Green Hermit, *Phaethornis guy* (PG); Brown Violetear, *Colibri delphinae* (CD); Black-throated Mango, *Anthracothorax nigricollis* (AN); White-vented Plumeleteer, *Chalybura buffonii* (CB); White-necked Jacobin, *Florisuga mellivora* (FM); Sparkling Violetear, *Colibri coruscans* (CC). Species' drawings provided by Fernando Ayerbe-Quiñones.

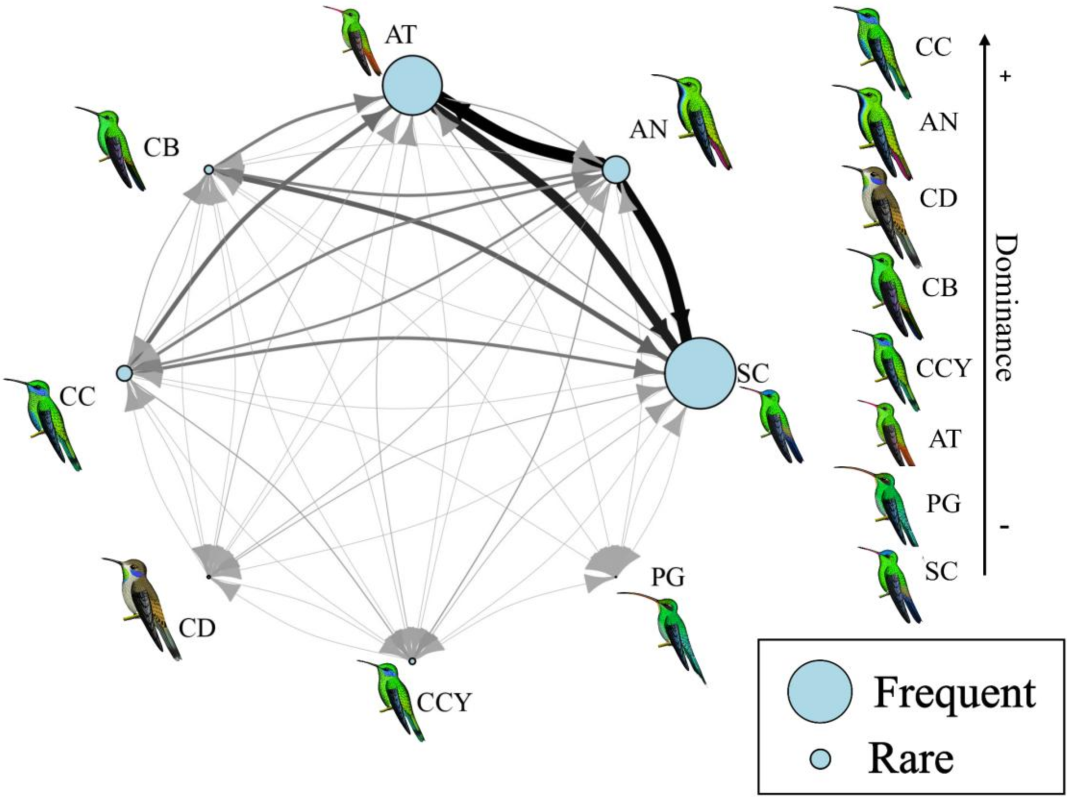


**Figure S1. Dominance hierarchy based on the total interaction matrix.** Interspecific interactions occurred primarily among *Colibri coruscans* (CC), *Chalybura buffonii* (CB), *Amazilia tzacatl* (AT), *Anthracothorax nigricollis* (AN) and *Saucerottia cyanifrons* (SC), with the latter three the most frequent visitors to the feeders. Larger and more aggressive species, such as *C. coruscans* (CC) and *A. nigricollis* (AN), occupied the highest ranks in the dominance hierarchy, while smaller but more frequent species, like *A. tzacatl* (AT) and *S. cyanifrons* (SC), ranked lower . *P. guy* (PG), a rare and non-territorial species, ranked at the bottom of the hierarchy.

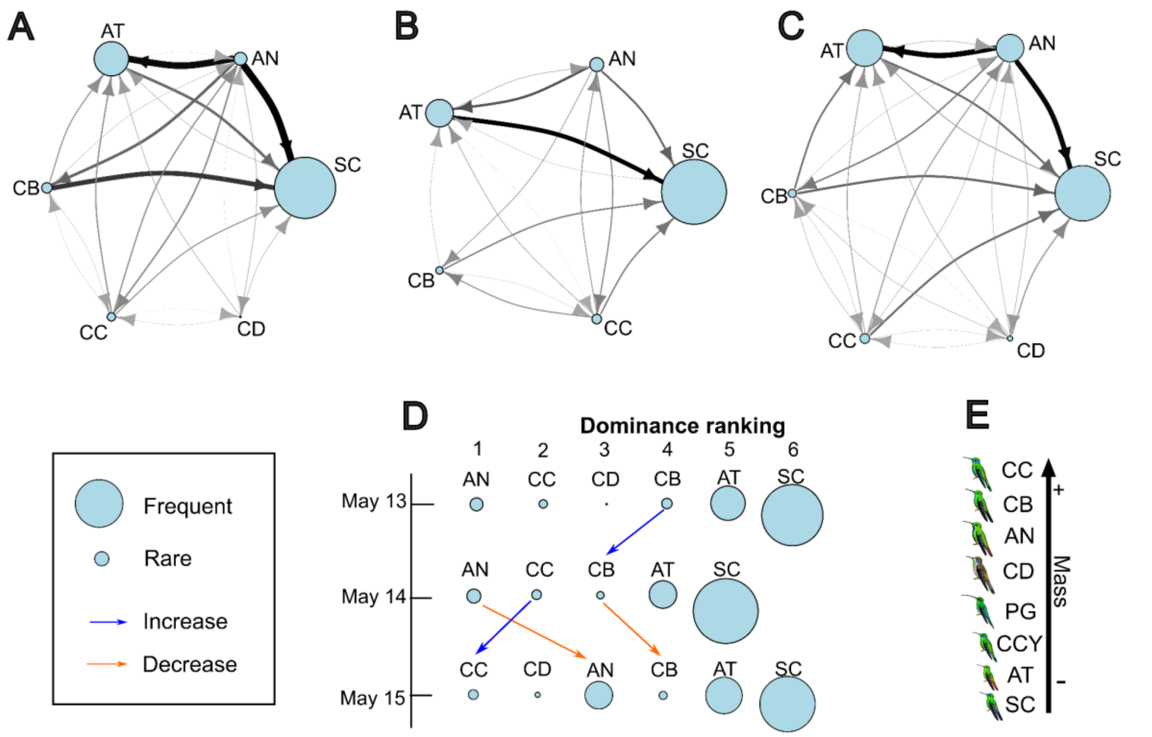


**Figure S2. Dominance variation across days in May.** A) May 13 network; B) May 14 network; C) May 15 network; D) Dominance ranking variation across days in May; E) Species' body mass scale. See Table 1 for the codes of hummingbird species. Dominance relationships were highly consistent between May 13 and 15, reflecting stable interspecific interactions during this month. *Anthracothorax nigricollis* (AN) showed a slight decrease in dominance on May 15, despite maintaining a consistent proportion of visits. *Colibri delphinae* (CD) was not recorded on May 14. *Amazilia tzacatl* (AT) and *Saucerottia cyanifrons* (SC) remained stable in both dominance rank and visit frequency throughout the days. Overall, network interconnectivity varied notably between days: On May 13, 2,359 aggressive interactions were recorded (8,287 visits); on May 14, 1,082 interactions (3,063 visits); and on May 15, 1,392 interactions (5,805 visits).

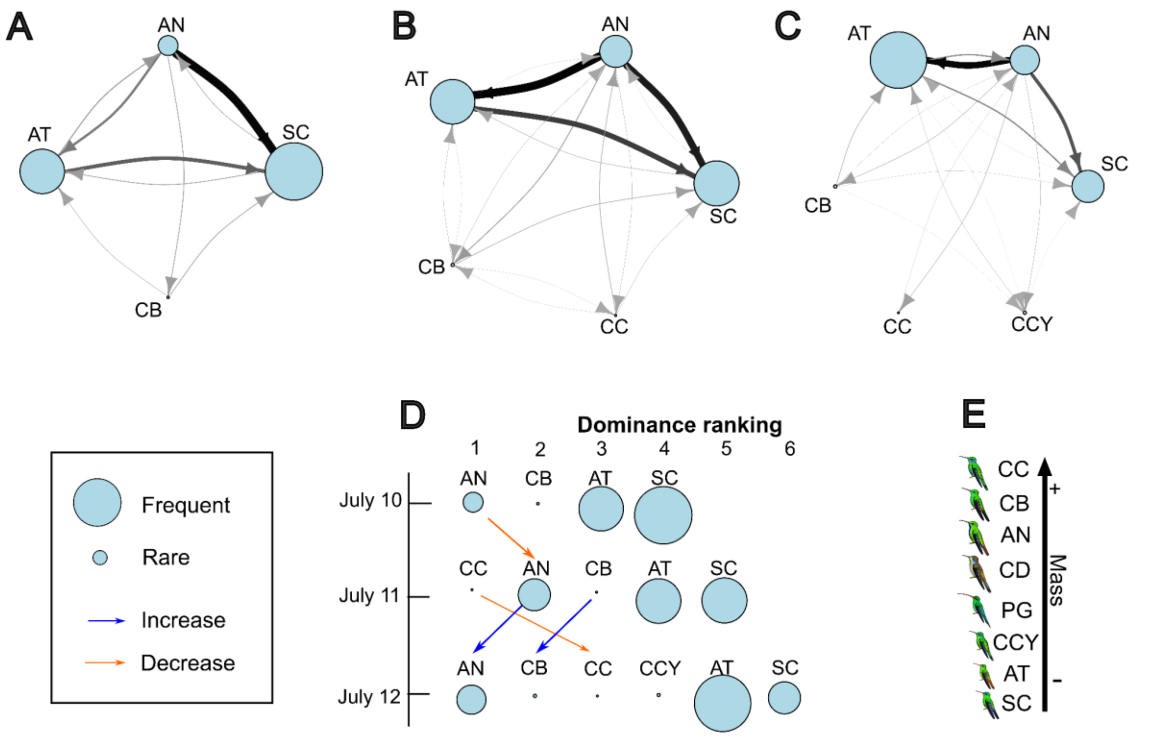


**Figure S3. Dominance variation across days in July.** A) July 10 network; B) July 11 network; C) July 12 network; D) Dominance ranking variation across days in July; E) Species' body mass scale. See Table 1 for the codes of hummingbird species. Among all months, July displayed the most dynamic shifts in dominance rankings across days. *Anthracothorax nigricollis* (AN) consistently held a dominant position with high visit frequency, whereas *Colibri coruscans* (CC) showed high variability and was absent from the interaction network on July 10, resulting in a notably sparse network that day. A total of 1,165 aggressive interactions were recorded on July 10 (3,722 visits), 1,745 on July 11 (5,313 visits), and 1,951 on July 12 (4,976 visits).

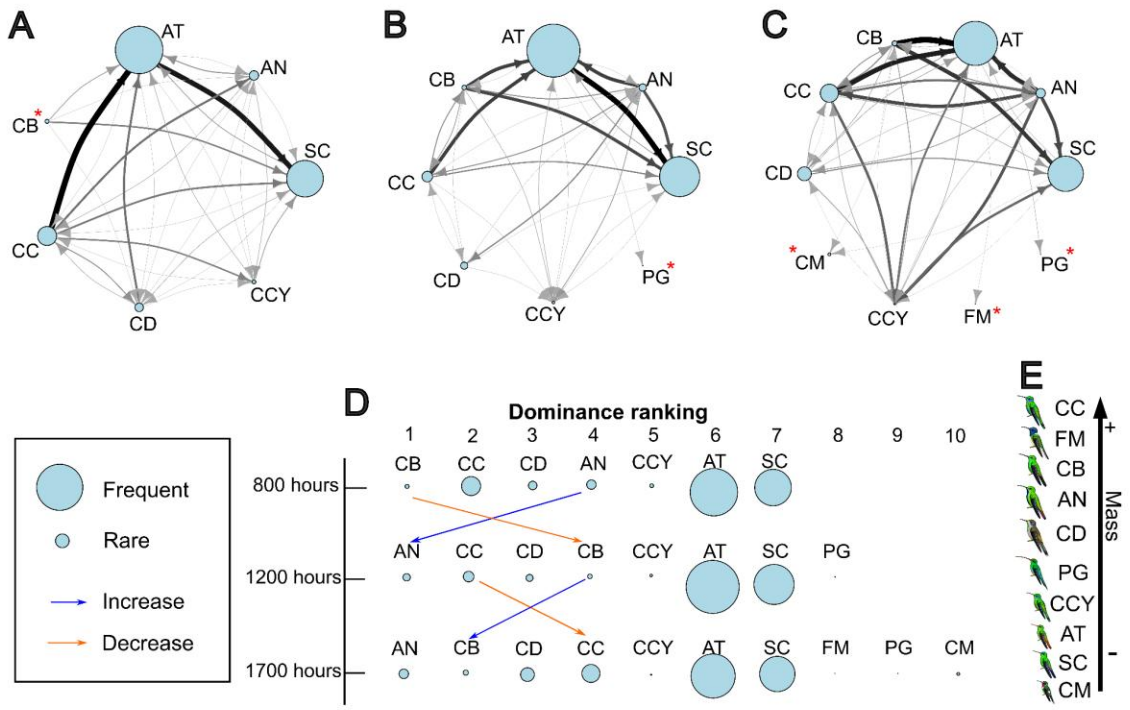


**Figure S4. Hourly variation in dominance variation on March 06.** A) March 06 - 0800 hours network; B) March 06 - 1200 hours network; C) March 06 - 1700 hours network; D) Dominance ranking variation on March 06; E) Species' body mass scale. See Table 1 for the codes of hummingbird species. On this day, interaction networks remained highly connected throughout the day. New species were observed in the late afternoon, particularly at 1700 hours. *Anthracothorax nigricollis* (AN) was not dominant during the early hour, while *Chalybura buffonii* (CB) exhibited notable fluctuations in dominance rank over the course of the day.

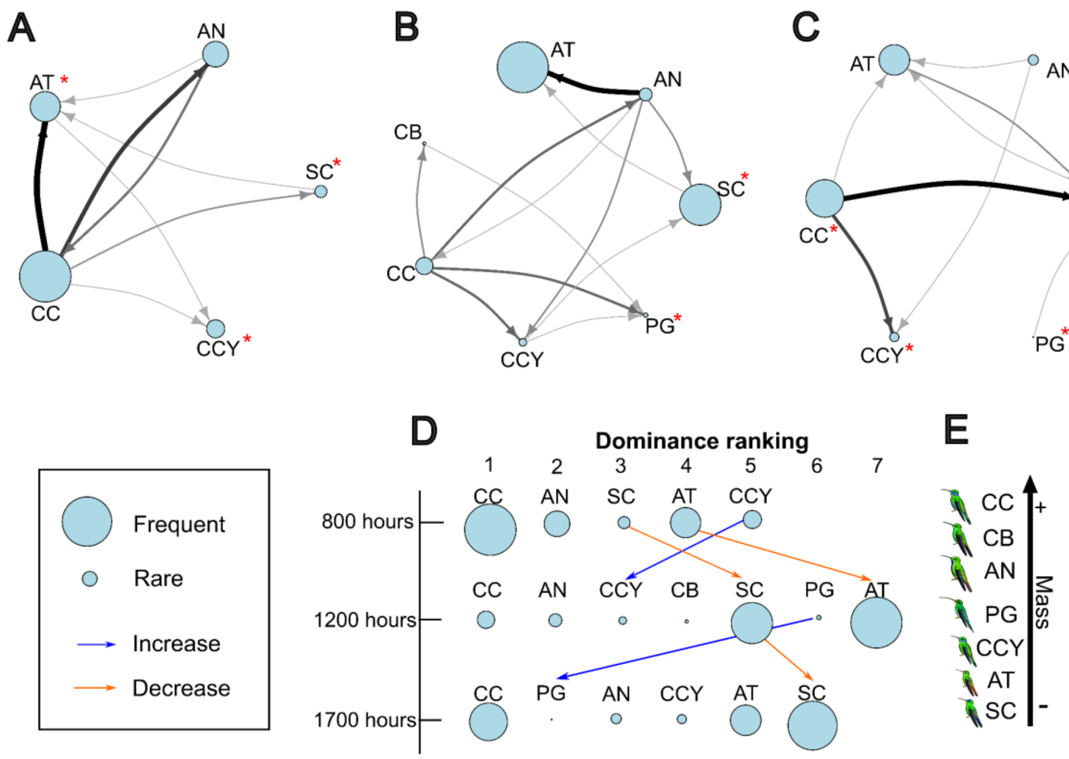


**Figure S5. Hourly variation in dominance variation on March 08.** A) March 08 - 0800 hours network; B) March 08 - 1200 hours network; C) March 08 - 1700 hours network; D) Dominance ranking variation on March 08; E) Species' body mass scale. See Table 1 for the codes of hummingbird species. On this day, *Colibri coruscans* (CC) maintained a dominant position throughout the day, regardless of its visit frequency. In contrast, *Saucerottia cyanifrons* (SC), typically a less dominant species, exhibited some variation in dominance across hours. By 1700 hours, the interaction network included fewer species, with unusual patterns such as *Phaethornis guy* (PG) appearing in a dominant position late in the afternoon.

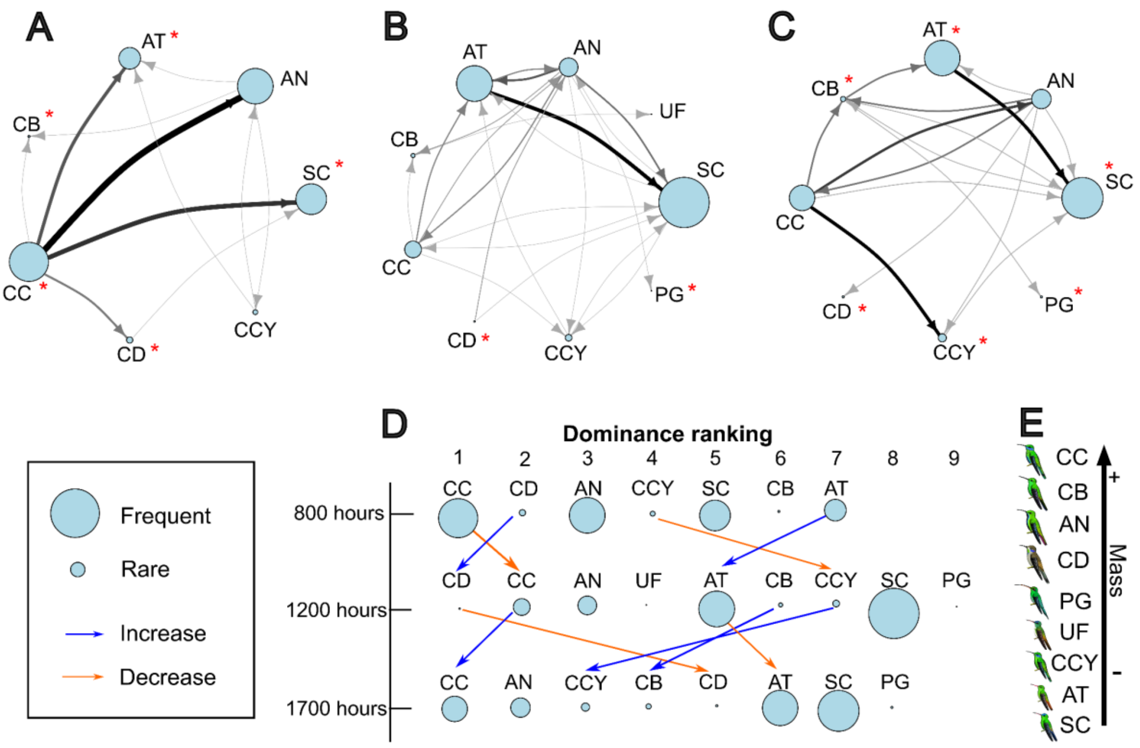


**Figure S6. Hourly variation in dominance on March 09.** A) March 09 - 0800 hours network; B) March 09 - 1200 hours network; C) March 09 - 1700 hours network; D) Dominance ranking variation on March 09; E) Species' body mass scale. See Table 1 for the codes of hummingbird species. This day showed greater variability in dominance ranks across all levels of the hierarchy. At the top level, *Colibri coruscans* (CC) dropped from the highest rank at 1200 hours, while *Colibri delphinae* (CD) declined at 1700 hours. In the mid-level ranks, *Colibri cyanotus* (CCY) showed positional shifts throughout the day, and *Amazilia tzacatl* (AT) ascended at 1200 hours before dropping again later. Additionally, new species—*Uranomitra franciae* (UF) and *Phaethornis guy* (PG)— entered the hierarchy at 1200 hours.

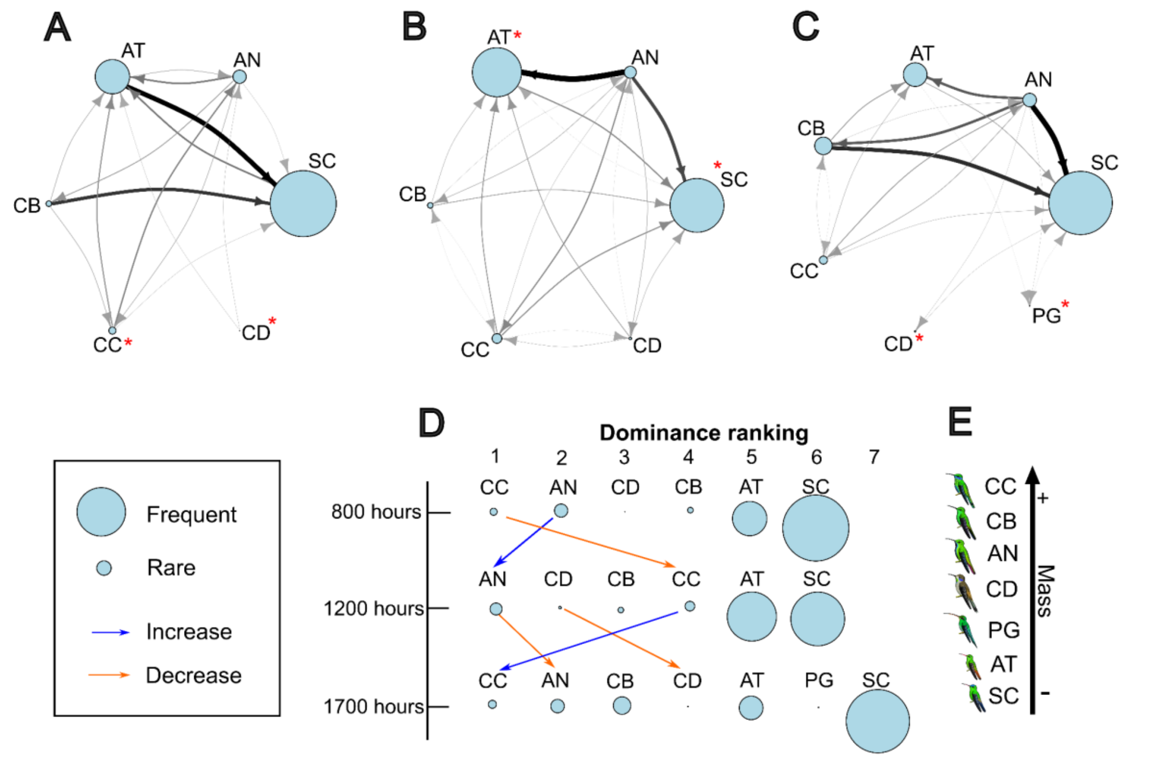


**Figure S7. Hourly variation in dominance on May 13.** A) May 13 - 0800 hours network; B) May 13 - 1200 hours network; C) May 13 - 1700 hours network; D) Dominance ranking variation on May 13; E) Species' body mass scale. See Table 1 for the codes of hummingbird species. *Colibri coruscans* (CC) and *Anthracothorax nigricollis* (AN) were dominant during the morning and late afternoon hours. However, at 1200 hours, *Colibri coruscans* (CC) experienced a notable drop in dominance. In contrast, *Amazilia tzacatl* (AT) and *Saucerottia cyanifrons* (SC) consistently occupied lower ranks in the hierarchy, in line with patterns observed across other temporal scales.

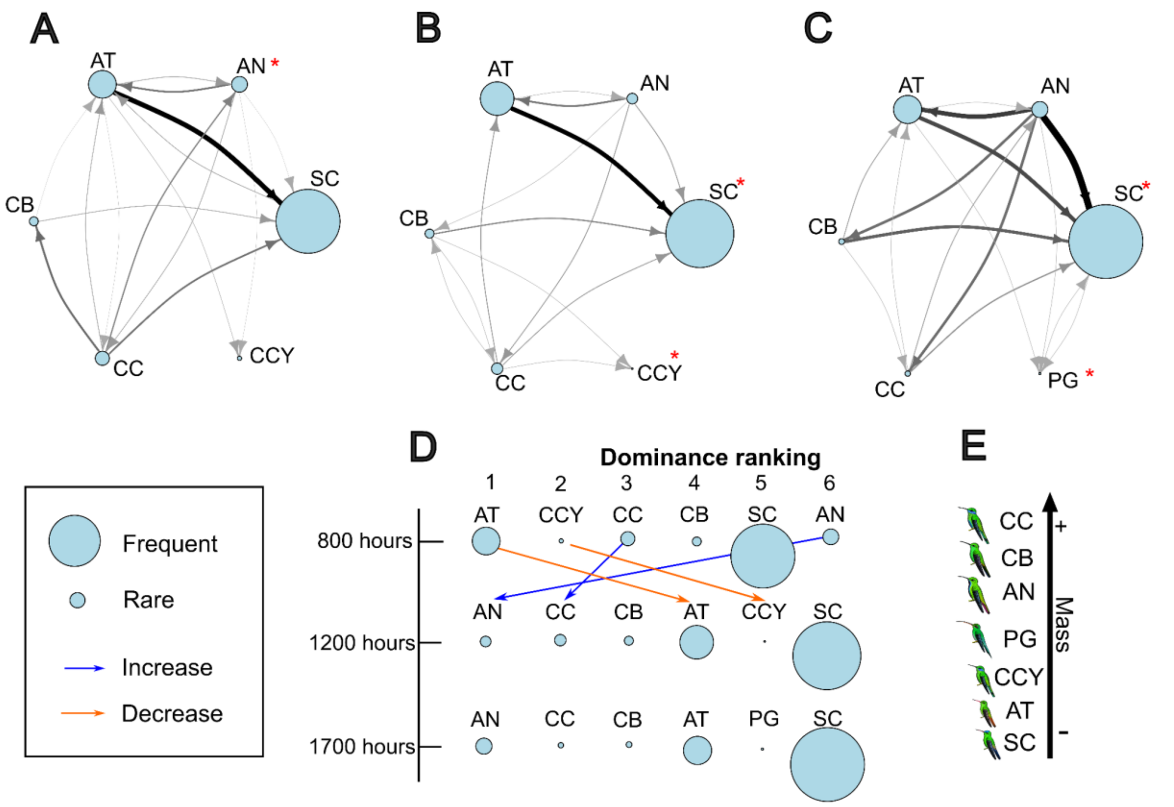


**Figure S8. Hourly variation in dominance on May 14.** A) May 14 - 0800 hours network; B) May 14 - 1200 hours network; C) May 14 - 1700 hours network; D) Dominance ranking variation on May 14; E) Species' body mass scale. See Table 1 for the codes of hummingbird species. An atypical dominance pattern was observed at 800 hours, with *Amazilia tzacatl* (AT) and *Colibri cyanotus* (CCY) occupying the top ranks, despite the presence of larger, typically more dominant species. *Anthracothorax nigricollis* (AN) held the lowest rank during this time but regained dominance for the remainder of the day.

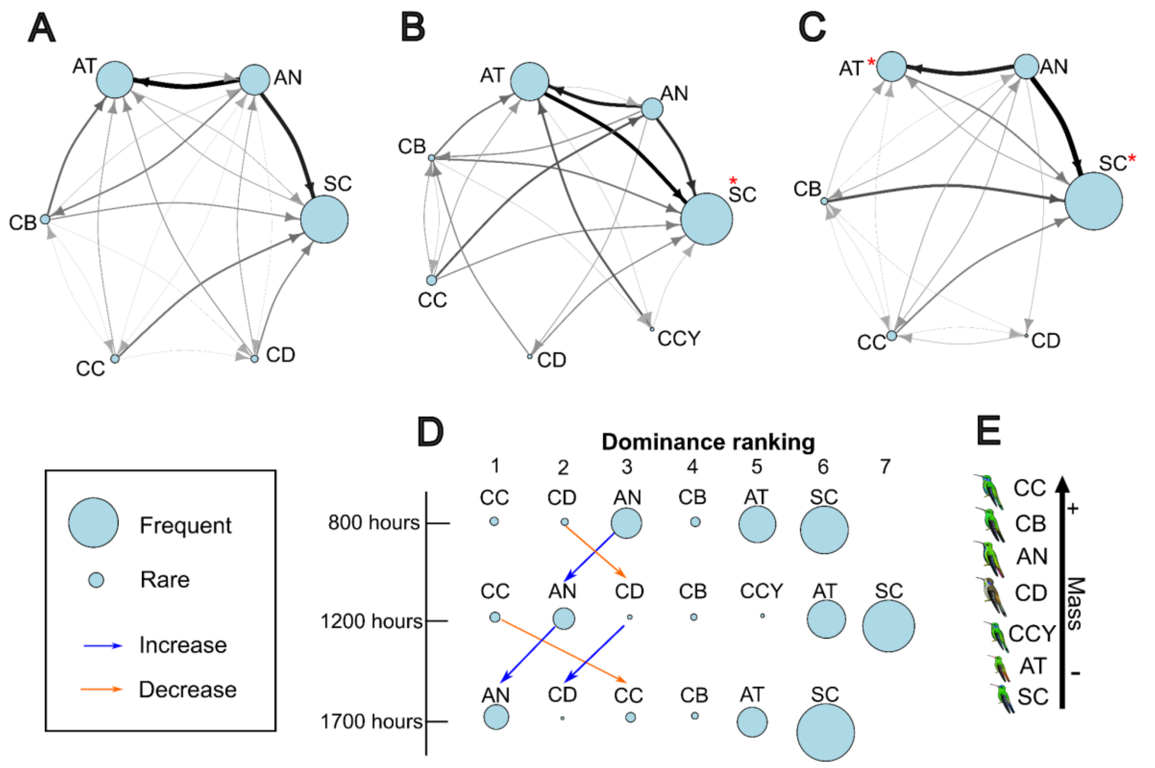


**Figure S9. Hourly variation in dominance on May 15.** A) May 15 - 0800 hours network; B) May 15 - 1200 hours network; C) May 15 - 1700 hours network; D) Dominance ranking variation on May 15; E) Species' body mass scale. See Table 1 for the codes of hummingbird species. Frequent shifts occurred among the top ranks, with *Anthracothorax nigricollis* (AN), *Colibri coruscans* (CC), and *Colibri delphinae* (CD) alternating in dominance throughout the day. Despite these fluctuations, their visit frequencies remained relatively stable. In contrast, mid-ranked positions showed minimal variation across hours.

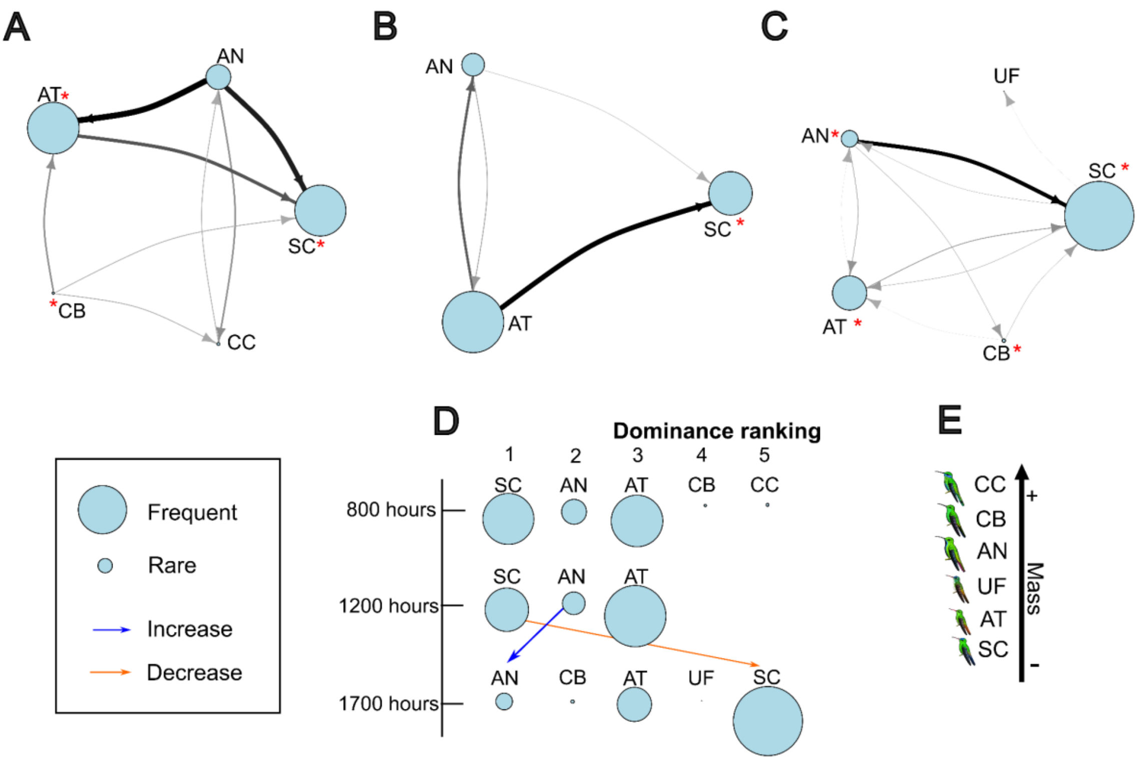


**Figure S10. Hourly variation in dominance on July 10.** A) July 10 - 0800 hours network; B) July 10 - 1200 hours network; C) July 10 - 1700 hours network; D) Dominance ranking variation on July 10; E) Species' body mass scale. See Table 1 for the codes of hummingbird species. At 800 hours, *Saucerottia cyanifrons* (SC) exhibited high dominance, while larger species such as *Chalybura buffonii* (CB) and *Colibri coruscans* (CC) occupied lower ranks. By 1200 hours, the dominance network had fewer species, with SC maintaining the top position. At 1700 hours, the network shifted to resembled patterns observed during other temporal periods.

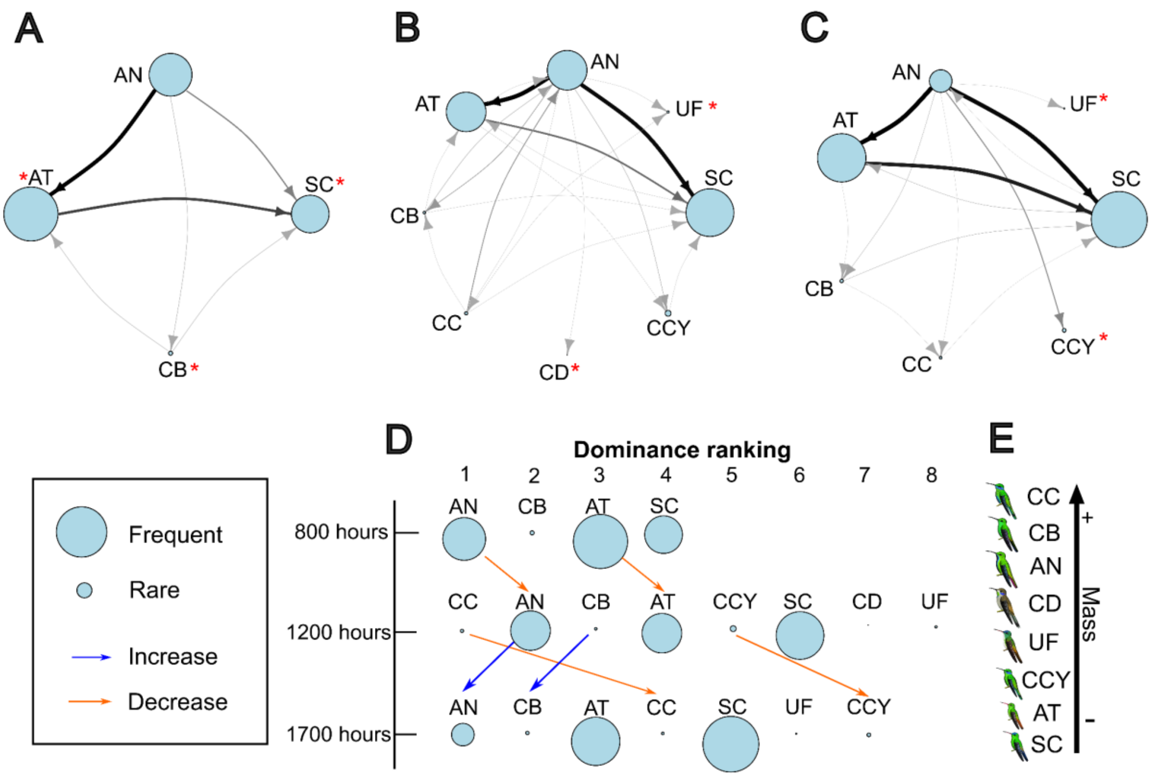


**Figure S11. Hourly variation in dominance on July 11.** A) July 11 - 0800 hours network; B) July 11 - 1200 hours network; C) July 11 - 1700 hours network; D) Dominance ranking variation on July 11; E) Species' body mass scale. See Table 1 for the codes of hummingbird species. Species diversity was lower at 800 hours compared to later hours. *Anthracothorax nigricollis* (AN) consistently maintained a dominant position throughout the day, while *Colibri coruscans* (CC) only reached a high rank at 1200 hours. Notably, *Amazilia tzacatl* (AT), which typically ranks at the bottom of the hierarchy, was observed occupying mid-level positions—an atypical pattern for this species.

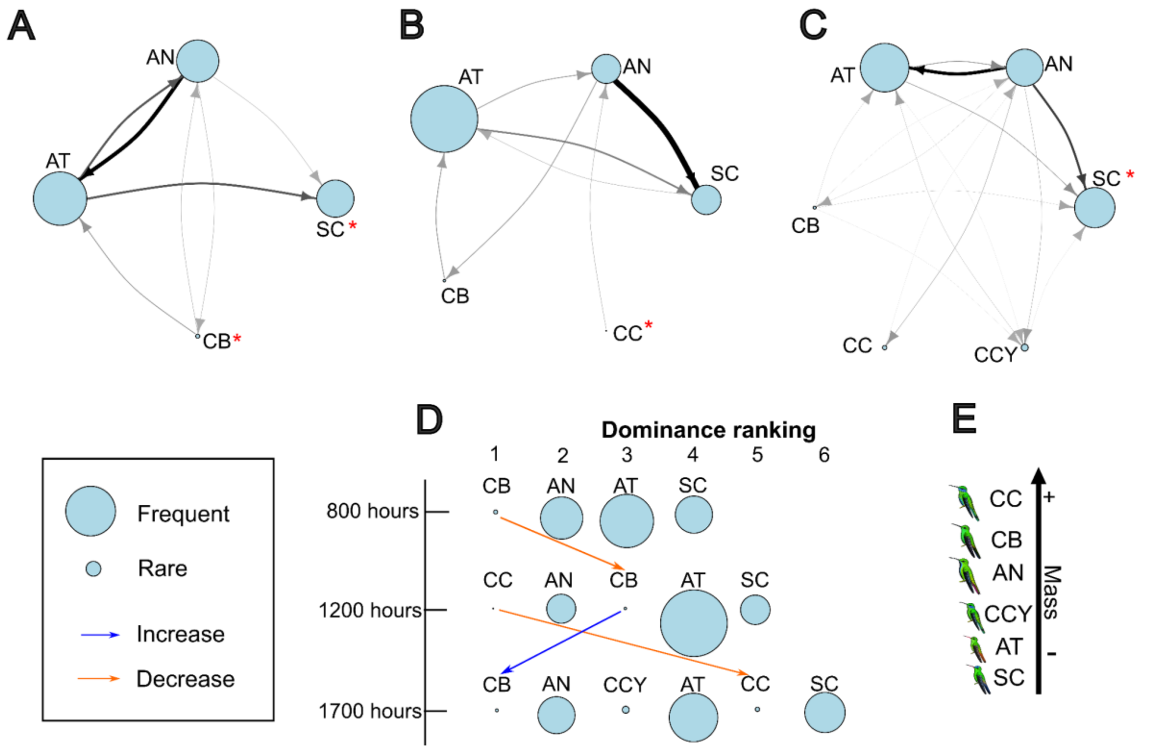


**Figure S12. Hourly variation in dominance on July 12.** A) July 12 - 0800 hours network; B) July 12 - 1200 hours network; C) July 12 - 1700 hours network; D) Dominance ranking variation on July 12; E) Species' body mass scale. See Table 1 for the codes of hummingbird species. *Chalybura buffonii* (CB) and *Anthracothorax nigricollis* (AN) were the most dominant species throughout the day. Similar to the previous day, *Colibri coruscans* (CC) ranked highly at 1200 hours but dropped significantly in later hours. *Amazilia tzacatl* (AT) and *Saucerottia cyanifrons* (SC) consistently remained in the lower ranks of the hierarchy. Overall, dominance certainty across species was higher on this day compared to other days in July.

**Table S2. Out-degree values per month and day.** Higher out-degree values indicate species initiating more aggressive interactions. In March, most species show relatively balanced interaction levels, while May exhibits the highest overall out-degree values, especially for *Anthracothorax nigricollis* (AN) and *Colibri coruscans* (CC). In July, *C. coruscans* (CC) showed fewer attacks compared to other dominant species such as *A. nigricollis* (AN) and *Chalybura buffonii* (CB). Daily variation is also observed, with SC consistently showing the lowest out-degree, while *A. nigricollis* (AN) and *C. coruscans* (CC) stand out on May 13. Species codes are provided in Table 1.

| Species | All data | March | 6-mar | 8-mar | 9-mar | May | 13-may | 14-may | 15-may | July | 10-jul | 11-jul | 12-jul |
|---|---|---|---|---|---|---|---|---|---|---|---|---|---|
| SC | 8 | 7 | 7 | 4 | 7 | 6 | 5 | 4 | 4 | 4 | 3 | 3 | 2 |
| AN | 8 | 7 | 7 | 7 | 7 | 7 | 6 | 5 | 7 | 6 | 5 | 6 | 5 |
| AT | 8 | 7 | 7 | 5 | 6 | 7 | 6 | 4 | 6 | 6 | 5 | 5 | 5 |
| CB | 8 | 7 | 7 | 6 | 7 | 7 | 6 | 6 | 7 | 6 | 4 | 6 | 4 |
| CC | 8 | 7 | 7 | 7 | 7 | 7 | 6 | 5 | 7 | 6 | 4 | 6 | 5 |
| CD | 7 | 7 | 7 | 4 | 5 | 6 | 6 | 3 | 6 | | | | |
| CCY | 8 | 7 | 7 | 4 | 5 | 6 | | | 5 | 5 | | 5 | |
| PG | 3 | | | | | | | | | | | | |

**Table S3. In-closeness values per month and day.** In-closeness reflects how quickly a species can be reached by others in the network. *Saucerottia cyanifrons* (SC) consistently record high in-closeness across all months, suggesting frequent interactions with many species. In May, *S. cyanifrons* (SC) and *Amazilia tzacatl* (AT) exhibit the highest in-closeness on May 13. On May 14 and 15, *S. cyanifrons* (SC) maintained the highest in-closeness values, reinforcing its central position despite its low dominance. Species codes are provided in Table 1.

| Species | All data | March | 6-mar | 8-mar | 9-mar | May | 13-may | 14-may | 15-may | July | 10-jul | 11-jul | 12-jul |
|---|---|---|---|---|---|---|---|---|---|---|---|---|---|
| SC | 1 | 1 | 1 | 0.857 | 0.667 | 1 | 1 | 1 | 1 | 1 | 1 | 1 | 1 |
| AN | 0.875 | 1 | 1 | 0.75 | 1 | 1 | 1 | 0.833 | 0.857 | 1 | 1 | 1 | 0.8 |
| AT | 1 | 1 | 1 | 1 | 0.75 | 1 | 1 | 1 | 1 | 1 | 1 | 1 | 1 |
| CB | 0.875 | 1 | 1 | 0.667 | 0.857 | 1 | 1 | 0.714 | 0.857 | 0.833 | 0.8 | 0.833 | 0.8 |
| CC | 0.857 | 1 | 1 | 0.857 | 1 | 1 | 1 | 0.833 | 0.857 | 0.714 | 0.667 | 0.625 | 0.667 |
| CD | 0.875 | 1 | 1 | 0.667 | 0.75 | 0.75 | 0.833 | 0.5 | 0.667 | | | | |
| CCY | 0.875 | 1 | 1 | 1 | 0.75 | 0.857 | | | 0.857 | 1 | | 0.833 | |
| PG | 0.875 | | | | | | | | | | | | |

**Table S4. Out-closeness values per month and day.** Out-closeness indicates how quickly a species can reach others in the network. May shows the highest values overall, particularly for *Anthracothorax nigricollis* (AN) and *Colibri coruscans* (CC). In contrast, *Saucerottia cyanifrons* (SC) and *Colibri cyanotus* (CCY) consistently exhibit the lowest out-closeness values, especially in July. Day-to-day variation is evident, with *S. cyanifrons* (SC) consistently low, while *A. nigricollis* (AN) and *C. coruscans* (CC) display peaks in influence on May 13 and 14. Species codes are provided in Table 1.

| Species | All data | March | 6-mar | 8-mar | 9-mar | May | 13-may | 14-may | 15-may | July | 10-jul | 11-jul | 12-jul |
|---|---|---|---|---|---|---|---|---|---|---|---|---|---|
| SC | 1 | 1 | 1 | 0.667 | 1 | 0.857 | 0.833 | 0.714 | 0.667 | 0.714 | 0.667 | 0.625 | 0.571 |
| AN | 1 | 1 | 1 | 1 | 1 | 1 | 1 | 0.833 | 1 | 1 | 1 | 1 | 1 |
| AT | 1 | 1 | 1 | 0.750 | 0.857 | 1 | 1 | 0.625 | 0.857 | 1 | 1 | 0.833 | 1 |
| CB | 1 | 1 | 1 | 0.857 | 1 | 1 | 1 | 1 | 1 | 1 | 0.800 | 1 | 0.800 |
| CC | 1 | 1 | 1 | 1 | 1 | 1 | 1 | 0.833 | 1 | 1 | 1 | 1 | 1 |
| CD | 0.875 | 1 | 1 | 0.750 | 0.857 | 0.857 | 1 | 0.714 | 0.857 | | | | |
| CCY | 1 | 1 | 1 | 0.750 | 0.750 | 0.857 | | | 0.750 | 0.833 | | 0.833 | |
| PG | 0.583 | | | | | | | | | | | | |

**Table S5. Network-level indices per month and day: density, reciprocity, average path length, and out-degree centralization.** March displays the highest network density and reciprocity, indicating more interconnected and mutual interactions. May shows the highest out-degree centralization and path length, suggesting a more hierarchical and less evenly connected network. These values reflect the fluctuating structure and complexity of hummingbird dominance interactions across time.

| Temporality | Density | Reciprocity | Average path length | Out-degree centralization |
|---|---|---|---|---|
| **All data** | 1 | 0.92 | 1.11 | 0.11 |
| **March** | 1 | 1 | 1 | 0 |
| **6-mar** | 1 | 1 | 1 | 0 |
| **8-mar** | 0.88 | 0.75 | 1.24 | 0.29 |
| **9-mar** | 1 | 0.89 | 1.10 | 0.12 |
| **May** | 1 | 0.97 | 1.07 | 0.07 |
| **13-may** | 1 | 0.97 | 1.03 | 0.03 |
| **14-may** | 0.90 | 0.73 | 1.30 | 0.30 |
| **15-may** | 1 | 0.86 | 1.17 | 0.17 |
| **July** | 1 | 0.89 | 1.10 | 0.10 |
| **10-jul** | 1 | 0.82 | 1.15 | 0.20 |
| **11-jul** | 1 | 0.80 | 1.17 | 0.17 |
| **12-jul** | 1 | 0.75 | 1.20 | 0.20 |

**Table S6. Betweenness index per species, per month and day.** This measure captures how often a species serves as a bridge in the interaction network. In March, betweenness values are generally low across species. In May and July, *Saucerottia cyanifrons* (SC), *Colibri coruscans* (CC), and *Colibri cyanotus* (CCY) frequently show the lowest betweenness, while *Anthracothorax nigricollis* (AN) stands out on July 10 with the highest score. These patterns highlight which species play central or peripheral roles in dominance relationships. Species codes are provided in Table 1.

| Species | All data | March | 6-mar | 8-mar | 9-mar | May | 13-may | 14-may | 15-may | July | 10-jul | 11-jul | 12-jul |
|---|---|---|---|---|---|---|---|---|---|---|---|---|---|
| SC | 0.063 | 0 | 0 | 0.008 | 0.032 | 0.007 | 0 | 0.075 | 0.015 | 0 | 0 | 0 | 0 |
| AN | 0.004 | 0 | 0 | 0.064 | 0.032 | 0.023 | 0.013 | 0.050 | 0.073 | 0.058 | 0.125 | 0.150 | 0.042 |
| AT | 0.063 | 0 | 0 | 0.067 | 0.025 | 0.023 | 0.013 | 0.033 | 0.037 | 0.058 | 0.125 | 0.050 | 0.292 |
| CB | 0.004 | 0 | 0 | 0.017 | 0.007 | 0.023 | 0.013 | 0.217 | 0.043 | 0.017 | 0 | 0.050 | 0 |
| CC | 0.004 | 0 | 0 | 0.117 | 0.025 | 0.023 | 0.013 | 0.075 | 0.043 | 0 | 0 | 0 | 0 |
| CD | 0 | 0 | 0 | 0.025 | 0.007 | 0 | 0 | 0 | 0 | | | | |
| CCY | 0.004 | 0 | 0 | 0.036 | 0.007 | 0 | | | 0.022 | 0.017 | | 0 | |
| PG | 0 | | | | | | | | | | | | |

**Table S7. Hubs index per species, per month and day.** Higher hub scores indicate species that often initiate interactions. *Anthracothorax nigricollis* (AN) and *Chalybura buffonii* (CB) consistently rank highest, especially on March 6 and May 14. In contrast, *Saucerottia cyanifrons* (SC) displays the lowest hub values across most time periods, reflecting a higher tendency to lose rather than initiate interactions. *Colibri coruscans* (CC) reaches peak hub scores on March 8 and May 13. Species codes are provided in Table 1.

| Species | All data | March | 6-mar | 8-mar | 9-mar | May | 13-may | 14-may | 15-may | July | 10-jul | 11-jul | 12-jul |
|---|---|---|---|---|---|---|---|---|---|---|---|---|---|
| SC | 1 | 1 | 1 | 0.657 | 1 | 0.889 | 0.854 | 0.775 | 0.606 | 0.716 | 0.699 | 0.561 | 0.445 |
| AN | 1 | 1 | 1 | 1 | 1 | 1 | 1 | 0.958 | 1 | 1 | 1 | 1 | 1 |
| AT | 1 | 1 | 1 | 0.729 | 0.865 | 1 | 1 | 0.802 | 0.898 | 1 | 1 | 0.895 | 1 |
| CB | 1 | 1 | 1 | 0.911 | 1 | 1 | 1 | 1 | 1 | 1 | 0.896 | 1 | 0.844 |
| CC | 1 | 1 | 1 | 1 | 1 | 1 | 1 | 0.958 | 1 | 1 | 0.896 | 1 | 1 |
| CD | 0.885 | 1 | 1 | 0.612 | 0.762 | 0.868 | 1 | 0.613 | 0.861 | | | | |
| CCY | 1 | 1 | 1 | 0.637 | 0.762 | 0.889 | | | 0.756 | 0.872 | | 0.895 | |
| PG | 0.378 | | | | | | | | | | | | |

**Table S8. Authorities index per species, per month and day.** Higher authority values suggest species frequently targeted by others or prone to losing interactions. *Saucerottia cyanifrons* (SC) consistently ranks highest in authority scores, particularly in May and July, while *Colibri coruscans* (CC) often exhibits the lowest values. These results emphasize contrasting dominance dynamics between species, with *S. cyanifrons* (SC) often on the receiving end of aggression despite its central network position. Species codes are provided in Table 1.

| Species | All data | March | 6-mar | 8-mar | 9-mar | May | 13-may | 14-may | 15-may | July | 10-jul | 11-jul | 12-jul |
|---|---|---|---|---|---|---|---|---|---|---|---|---|---|
| SC | 1 | 1 | 1 | 0,890 | 1 | 1 | 1 | 1 | 1 | 1 | 1 | 1 | 1 |
| AN | 0,948 | 1 | 1 | 0,771 | 1 | 1 | 1 | 0,880 | 0,877 | 1 | 1 | 1 | 0,896 |
| AT | 1 | 1 | 1 | 1 | 1 | 1 | 1 | 1 | 1 | 1 | 1 | 1 | 1 |
| CB | 0,948 | 1 | 1 | 0,635 | 0,762 | 1 | 1 | 0,723 | 0,901 | 0,872 | 0,844 | 0,895 | 0,896 |
| CC | 0,948 | 1 | 1 | 0,869 | 0,865 | 1 | 1 | 0,848 | 0,901 | 0,716 | 0,445 | 0,561 | 0,699 |
| CD | 0,948 | 1 | 1 | 0,492 | 0,762 | 0,732 | 0,854 | 0,196 | 0,631 | | | | |
| CCY | 0,948 | 1 | 1 | 0,885 | 1 | 0,869 | | | 0,859 | 1 | | 0,895 | |
| PG | 0,878 | | | | | | | | | | | | |

**Table S9. Network densities of species interactions per site and month based on presence/absence matrices.** All networks show densities significantly lower than 1, indicating non-random, specific patterns of interactions among species. Exceptions include March, July, and the specific days of March 6 and 9, where densities were not significantly different from 1.

| | All data | March | March | | | May | May | | | July | July | | |
|---|---|---|---|---|---|---|---|---|---|---|---|---|---|
| | | | 6-mar | 8-mar | 9-mar | | 13-may | 14-may | 15-may | | 10-jul | 11-jul | 12-jul |
| **Density** | 0.893 | 1.000 | 1.000 | 0.762 | 0.905 | 0.929 | 0.967 | 0.733 | 0.833 | 0.900 | 0.850 | 0.833 | 0.800 |
| **Z-value** | NA | NA | NA | -2.397 | -1.827 | -1.833 | NA | -3.623 | -2.032 | -1.245 | -1.805 | -1.766 | -1.679 |
| **p-value** | 0 | 1.000 | 1.000 | 0.012 | 0.062 | 0.049 | 0 | 0.001 | 0.040 | 0.152 | 0.060 | 0.052 | 0.055 |

**Table S10. Comparison of presence/absence network densities across months and days within each site.** March 6 was the only day in March with a significantly different density compared to the other two days. Similarly, May 13 showed significantly different network density relative to May 14 and 15.

**Months**

| | May | July |
|---|---|---|
| **March** | $t$ = 0.3446, P = 0.3419 | $t$ = 0.1523, P = 0.4742 |
| **May** | - | $t$ = -0.0230, P = 0.5530 |

**March**

| | 08 March | 09 March |
|---|---|---|
| **06 March** | $t$ = 1.9111, P = 0.0448 | $t$ = 2.2072, P = 0.0296 |
| **08 March** | - | $t$ = 0.7794, P = 0.1935 |

**May**

| | 14 May | 15 May |
|---|---|---|
| **13 May** | $t$ = 2.2334, P = 0.0226 | $t$ = 2.7649, P = 0.0063 |
| **14 May** | - | $t$ = -1.0500, P = 0.8586 |

**July**

| | 11 July | 12 July |
|---|---|---|
| **10 July** | $t$ = -0.4592, P = 0.3034 | $t$ = -0.4970, P = 0.2684 |
| **11 July** | - | $t$ = -0.4736, P = 0.2594 |

**Table S11. Quadratic Assignment Procedure (QAP) correlations and corresponding *P*-values comparing species interaction networks across months.** Statistically significant *P*-values are indicated with **. A significant *P*-value indicates that the structure of interactions between two months is more similar than expected by chance. For example, if March–May and March–July show significant values, it suggests that the pattern of species interactions was conserved across those months.

| cor/p value | March | May | July |
|---|---|---|---|
| **March** | - | 0.006** | 0.02** |
| **May** | 0.61 | - | 0.892 |
| **July** | 0.462 | 0 | - |

**Table S12. Pearson correlations between species interaction networks across days and months, calculated using Quadratic Assignment Procedure (QAP).** Significance levels are denoted as follows: *:P<0.05, **: P<0.01, ***:P<0.001. Significant correlations between July–March, July–May, and May–March indicate that species maintained similar dominance interaction patterns across different months. Additionally, all day-to-day comparisons within the same month were significant, suggesting temporal consistency in dominance structure at shorter timescales.

**Months**

| | July | March | May |
|---|---|---|---|
| **July** | 1 | 0.523 ** | 0.866 ** |
| **March** | 0.523 ** | 1 | 0.688 ** |
| **May** | 0.866 ** | 0.688 ** | 1 |

**March**

| | 6-mar | 8-mar | 9-mar |
|---|---|---|---|
| **6-mar** | 1 | 0.502** | 0.72*** |
| **8-mar** | 0.502** | 1 | 0.647** |
| **9-mar** | 0.72*** | 0.647** | 1 |

**May**

| | 13-may | 14-may | 15-may |
|---|---|---|---|
| **13-may** | 1 | 0.737*** | 0.937*** |
| **14-may** | 0.737*** | 1 | 0.822*** |
| **15-may** | 0.937*** | 0.822*** | 1 |

**July**

| | 10-jul | 11-jul | 12-jul |
|---|---|---|---|
| **10-jul** | 1 | 0.894*** | 0.644** |
| **11-jul** | 0.894*** | 1 | 0.889** |
| **12-jul** | 0.644** | 0.889** | 1 |

**Table S13. Quadratic assignment procedure (QAP) correlations and *P*-values comparing species interaction networks across days within each month**. Significant correlations indicate structural similarity in interaction patterns between specific days. In March, significant correlations were found between March 9 and both March 6 and March 8. In May, all pairwise comparisons were significant: May 14 with May 13, May 15 with May 13, and May 15 with May 14. In July, significant correlations occurred between July 10 and 11, and between July 12 and 11.

| cor/p value | 6-mar | 8-mar | 9-mar |
|---|---|---|---|
| 6-mar | - | 0.066 | 0.003** |
| 8-mar | 0.309 | - | 0.018** |
| 9-mar | 0.559 | 0.661 | - |

| cor/p value | 13-may | 14-may | 15-may |
|---|---|---|---|
| 13-may | - | 0.023** | 0.007** |
| 14-may | 0.636 | - | 0.018** |
| 15-may | 0.944 | 0.646 | - |

| cor/p value | 10-jul | 11-jul | 12-jul |
|---|---|---|---|
| 10-jul | - | 0.007** | 0.057 |
| 11-jul | 0.823 | - | 0.009** |
| 12-jul | 0.645 | 0.912 | - |

**Conventions Figures S13 to S25.** Dominance certainty was higher in the species of lower rank; note the value of dominance certainty on the left side of the image and the dominance probability heatmap on the right, which depicts the simulated probability of being ranked subordinate or dominant based on our data (with the *x* axis showing losing species, and the *y* axis showing winning species). The heatmap's color scale signifies clearly defined relationships in which a species is linearly ranked as subordinate (blue region) or dominant (orange–red region), with the exception of when there were no data (white, 0.0, signifying intraspecific interactions), or when the dominance rank was unclear (brown region, ~0.5). Species' drawings provided by Fernando Ayerbe-Quiñones.

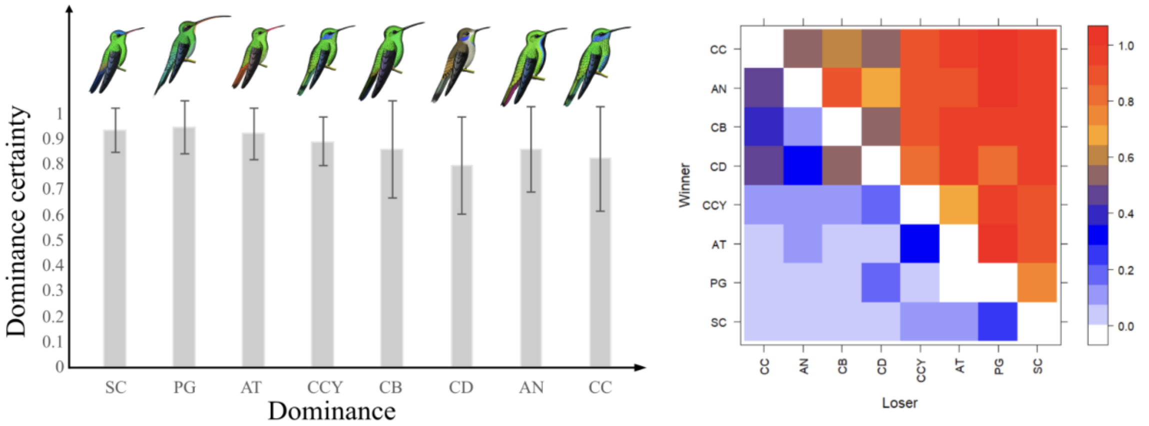


**Figure S13. Dominance certainty across the entire period.** Hummingbird species codes are provided in Table 1. Subordinate species generally exhibit the highest dominance certainty. In contrast, species like *Colibri delphinae* (CD) and *Colibri coruscans* (CC) show the lowest certainty, as indicated by the brown squares on the heatmap.

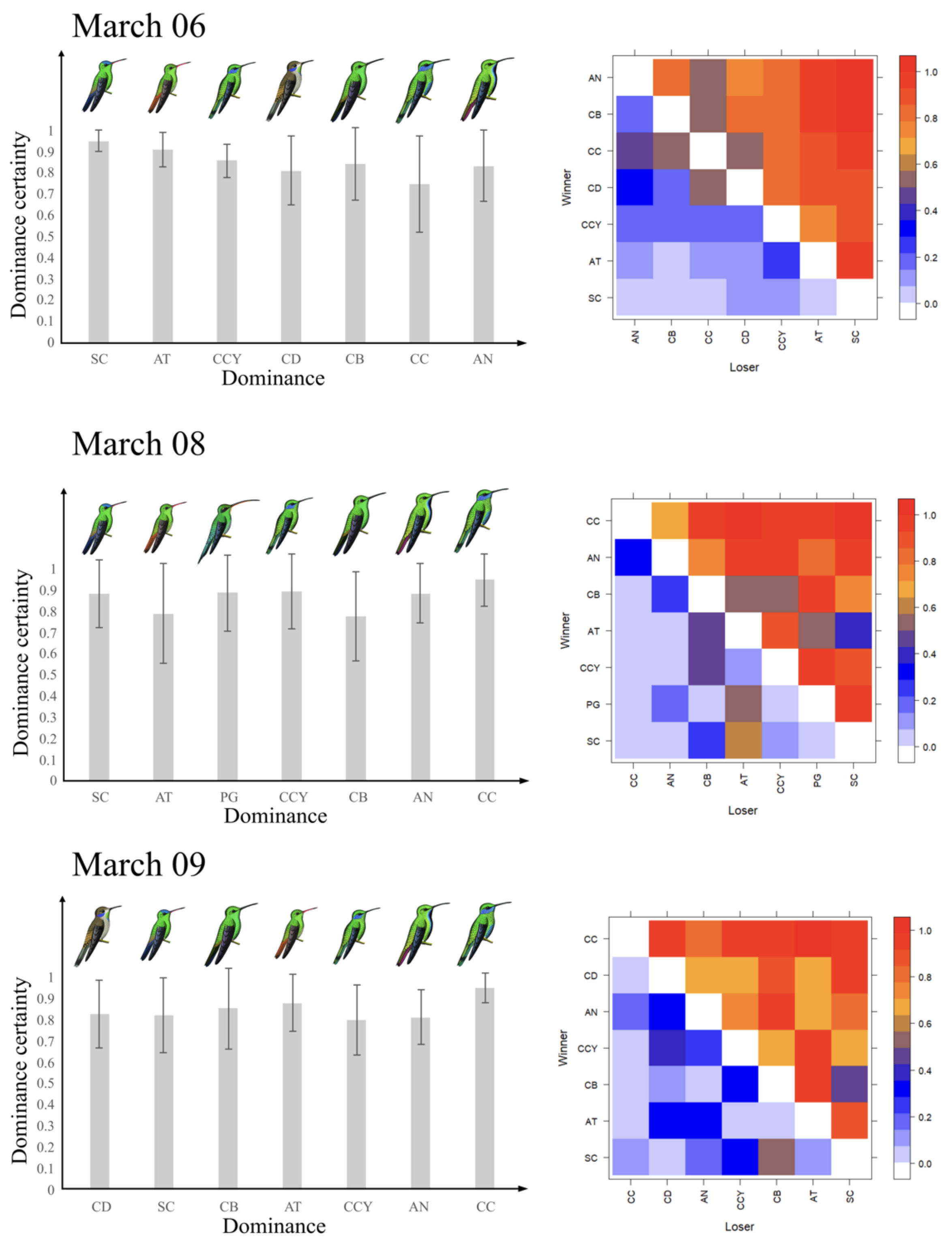


**Figure S14. Dominance certainty across days in March.** Species codes are listed in Table 1. Dominance certainty varied by species and day. On March 6, lower-ranking species showed the highest certainty, while on March 9, dominant species exhibited more consistent values. Species occupying intermediate ranks displayed high variability in dominance certainty throughout the three days.

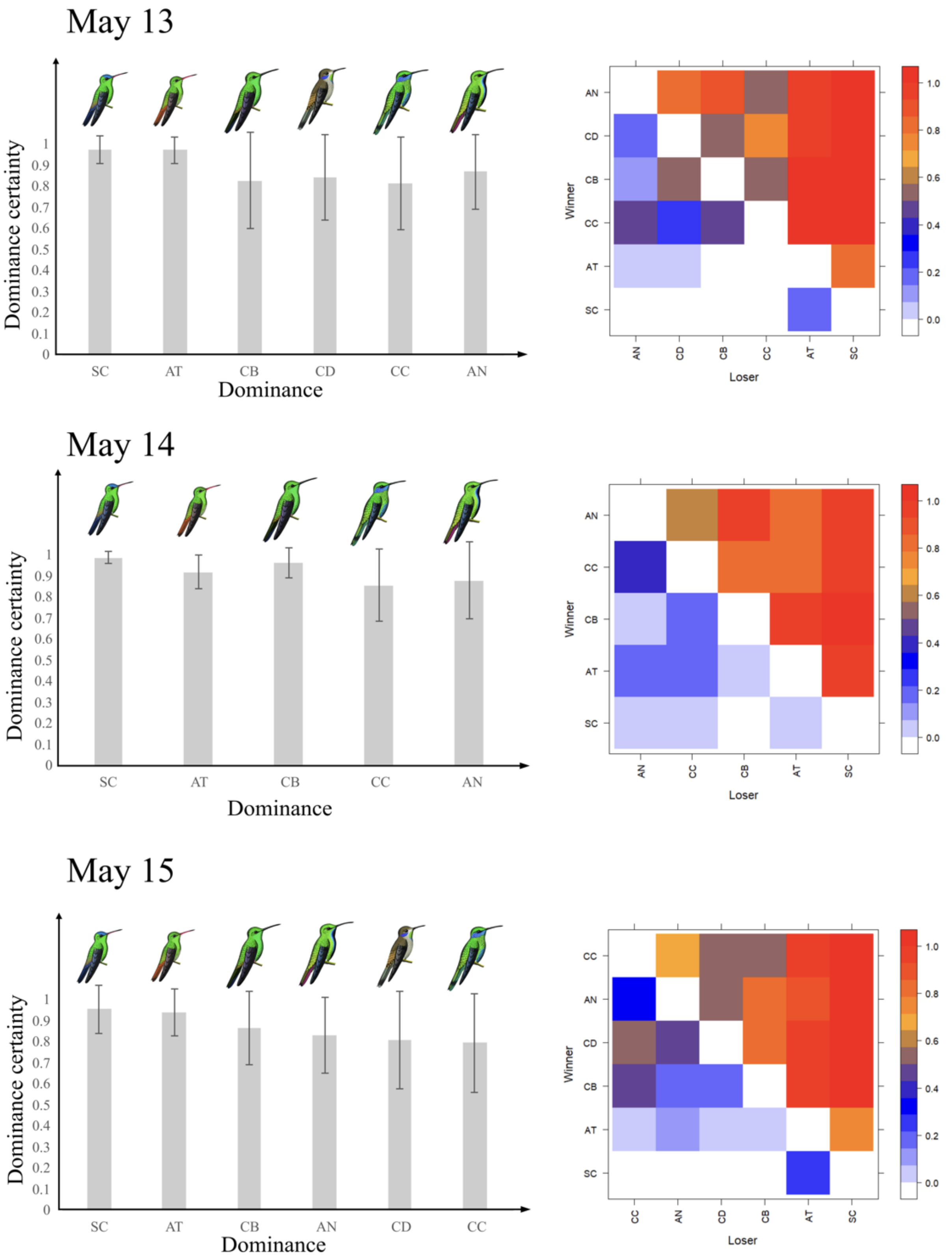


**Figure S15. Dominance certainty across days in May.** Species codes are listed in Table 1. In May, lower-ranking species such as *Amazilia tzacatl* (AT) and *Saucerottia cyanifrons* (SC) consistently exhibited higher dominance certainty compared to more dominant species.

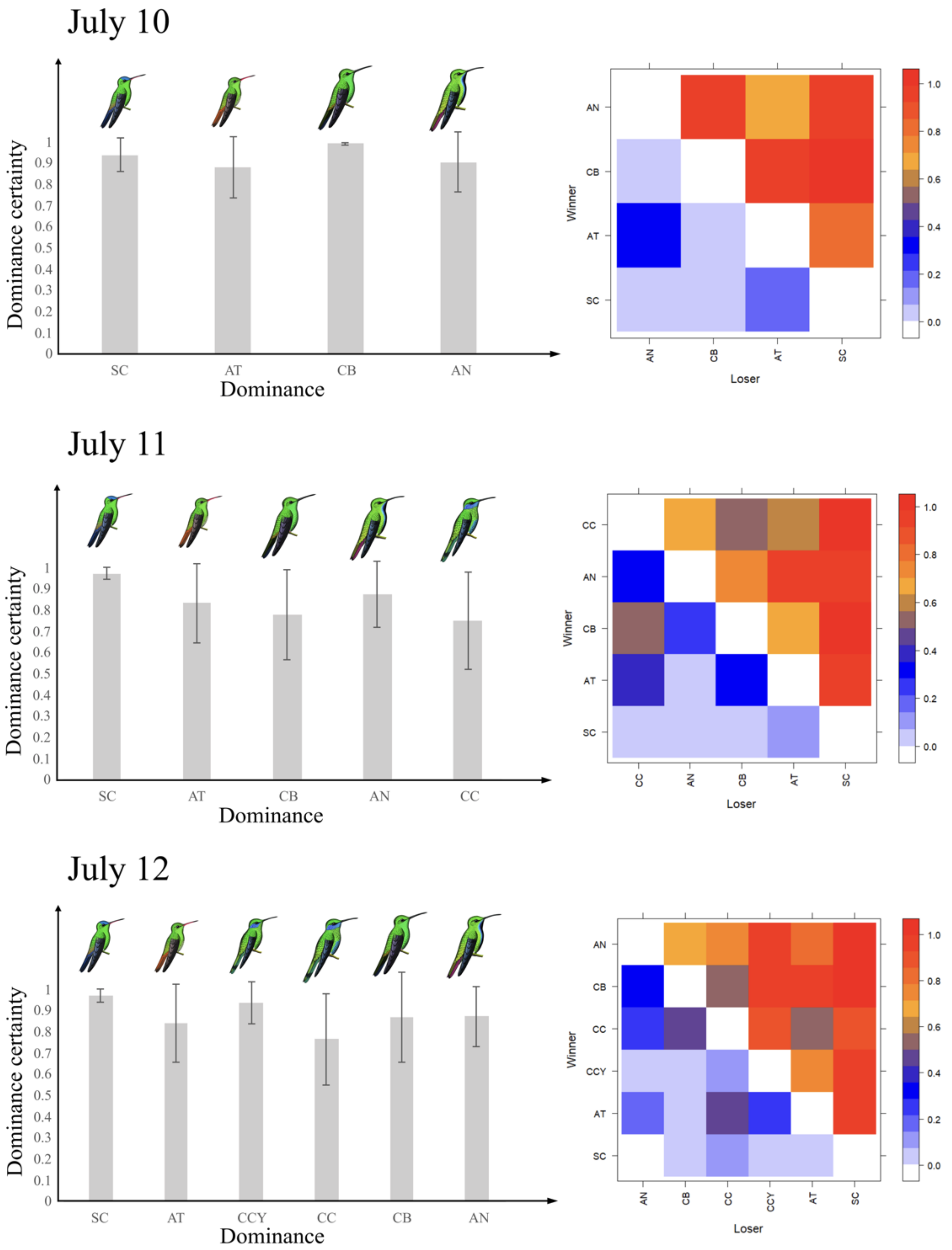


**Figure S16. Dominance certainty across days in July.** Species codes are listed in Table 1. Dominance certainty varied widely during July and was generally low across species. Notable exceptions included *Chalybura buffonii* (CB), which showed high certainty at 800 hours, and *Saucerottia cyanifrons* (SC), which consistently maintained high certainty throughout the day.

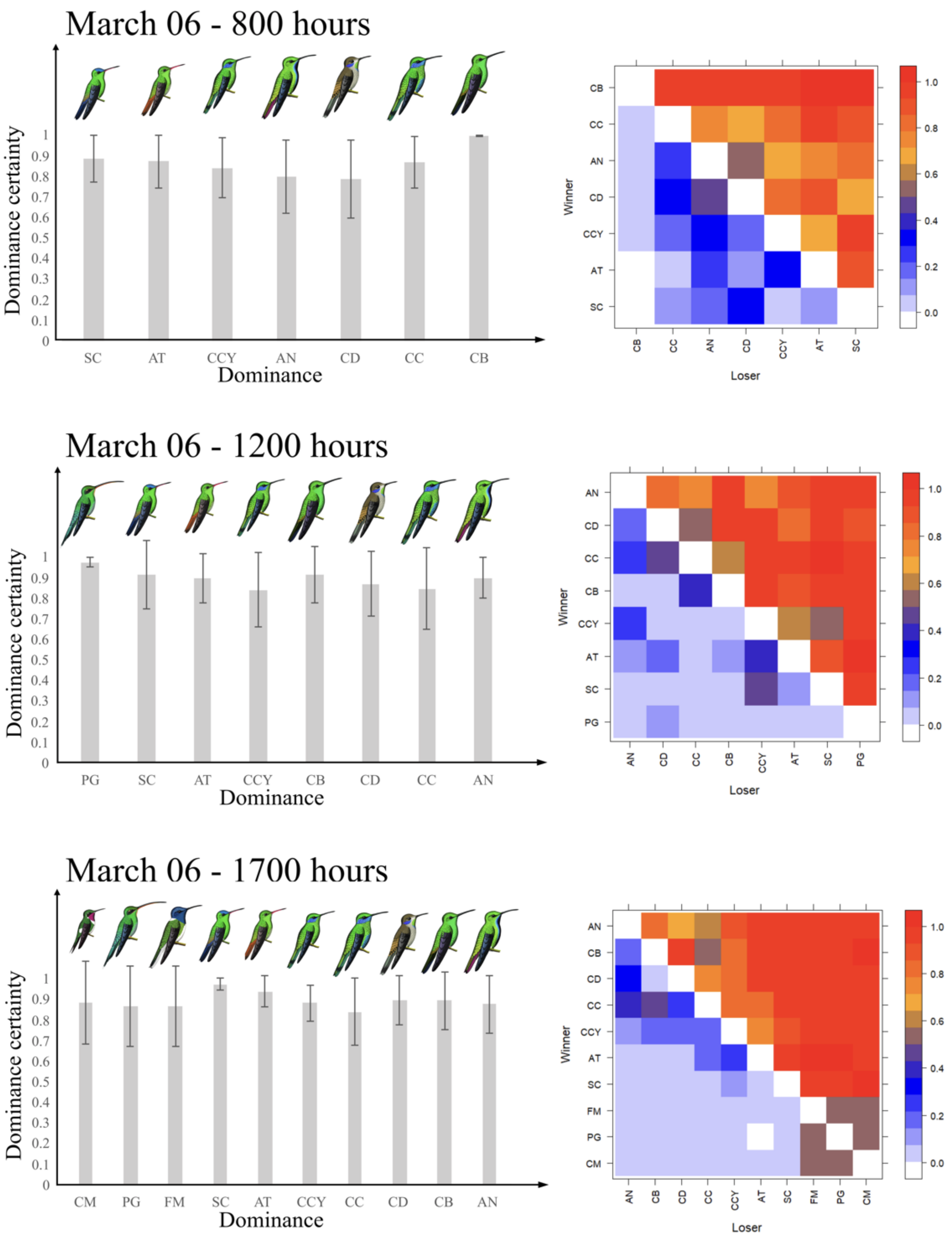


**Figure S17. Dominance certainty across hours on March 6.** Species codes are listed in Table 1. At 800 and 1200 hours, species at the top and bottom of the dominance hierarchy exhibited the highest certainty, while mid-ranked species showed greater variability. By 1700 hours, this pattern reversed, with lower certainty observed among species at both extremes of the hierarchy.

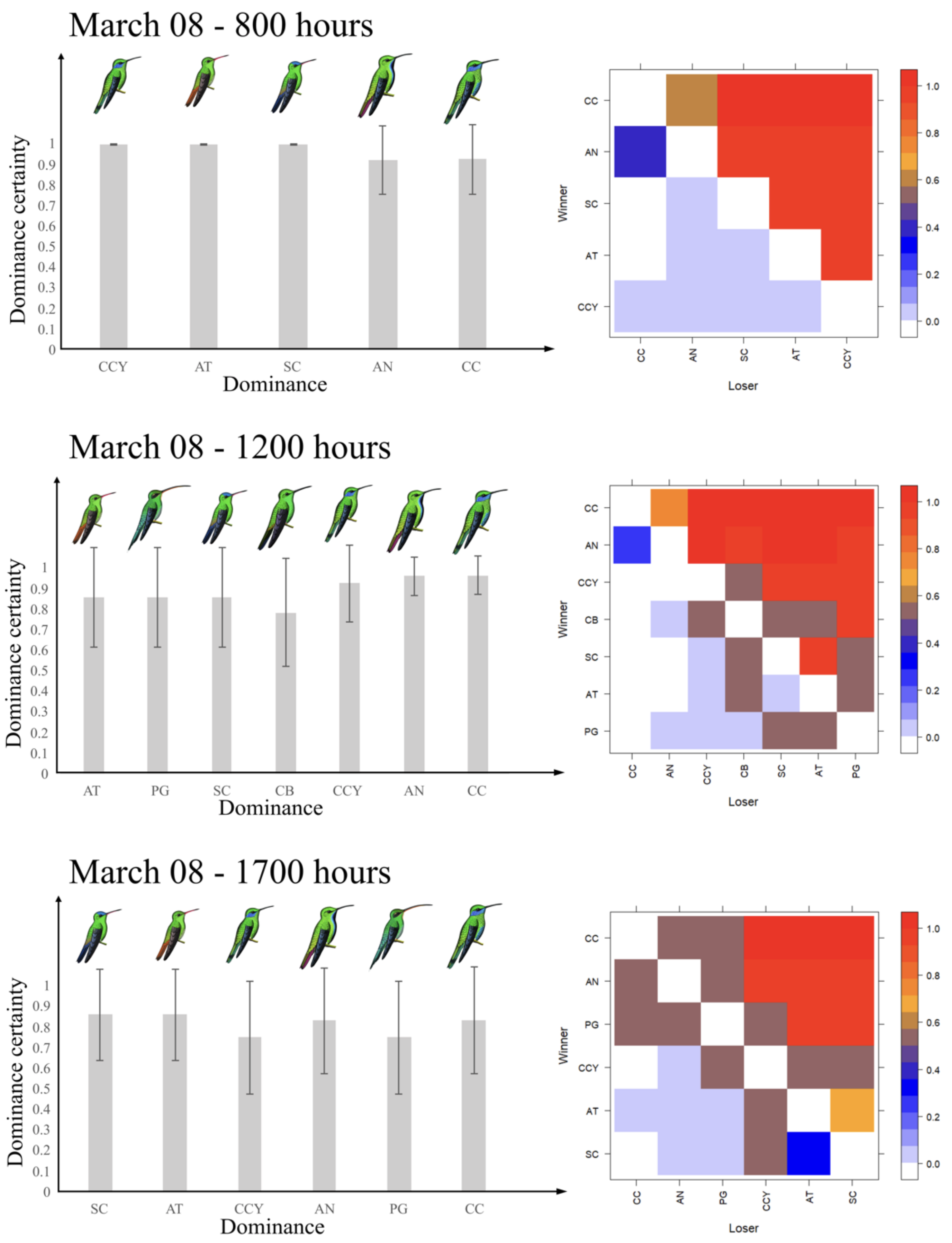


**Figure S18. Dominance certainty across hours on March 8.** Species codes are listed in Table 1. Overall dominance certainty was low throughout the day, with the exception of 800 hours, when species exhibited relatively higher certainty in their dominance ranks.

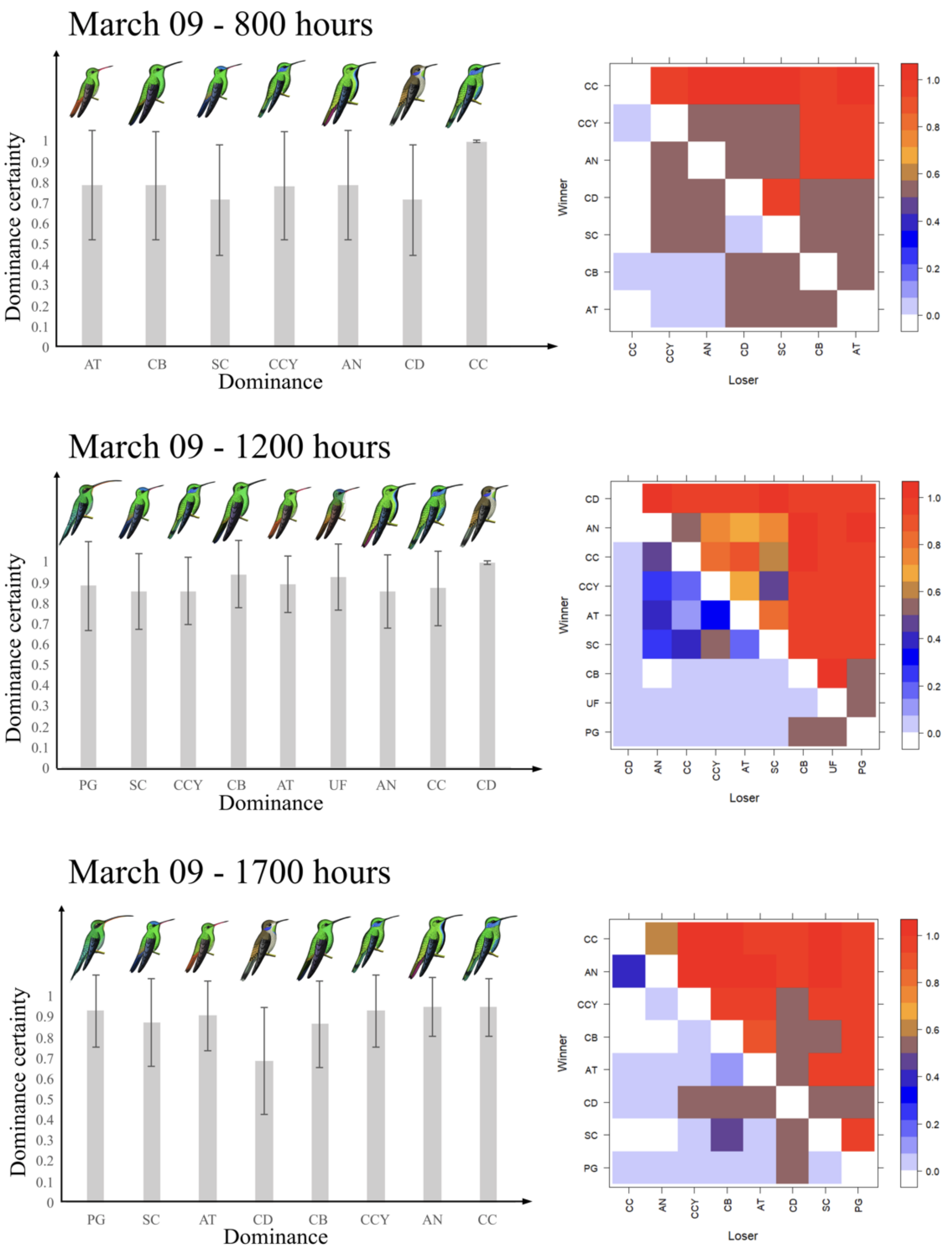


**Figure S19. Dominance certainty across hours on March 9.** Species codes are listed in Table 1. Overall dominance certainty was low throughout the day. Subordinate species consistently exhibited low certainty, whereas dominant species showed relatively higher and more stable certainty in their dominance relationships.

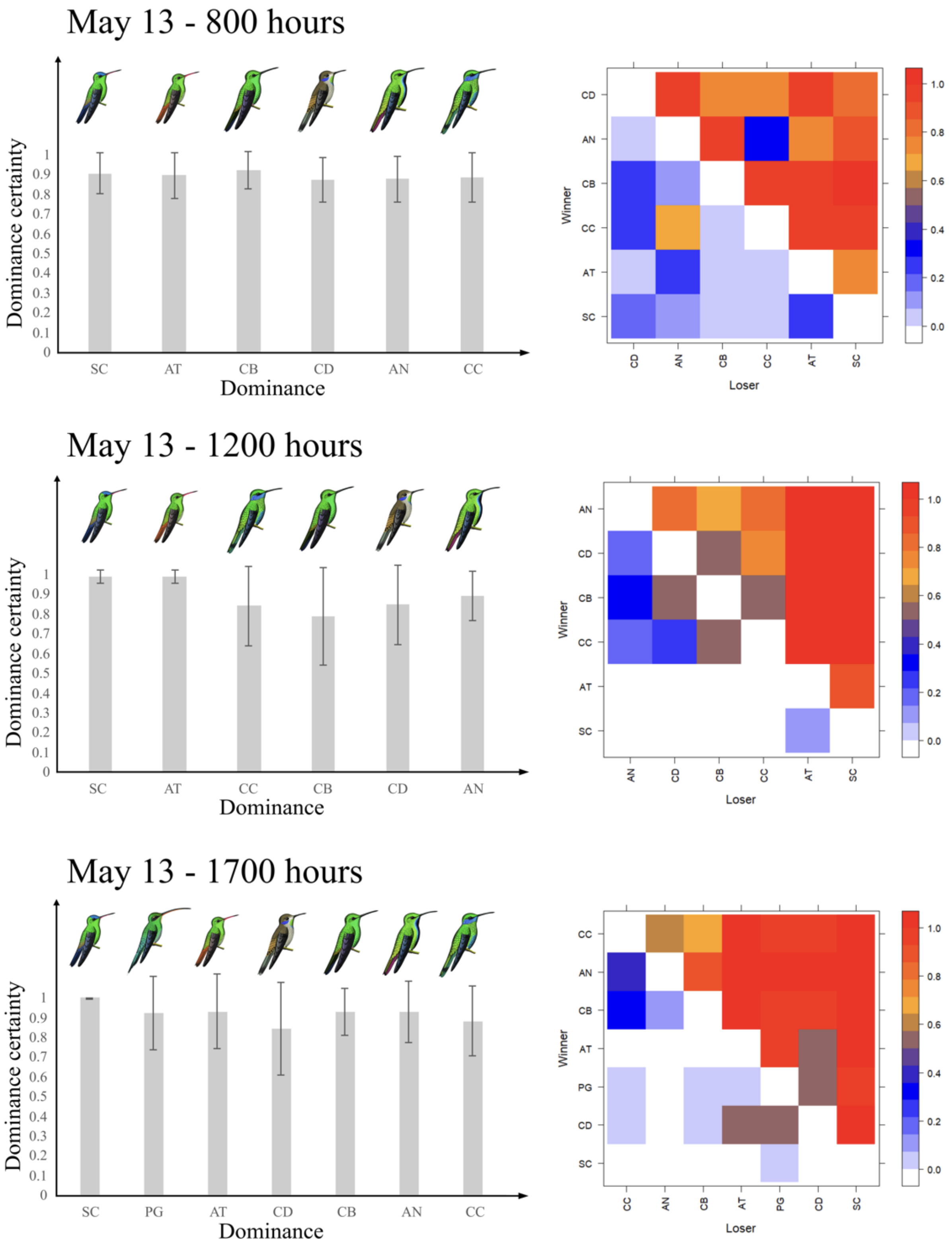


**Figure S20. Dominance certainty across hours on May 13.** Species codes are listed in Table 1. Dominance certainty was highest at 800 hours but declined over the course of the day, with the greatest variability observed at 1200 hours.

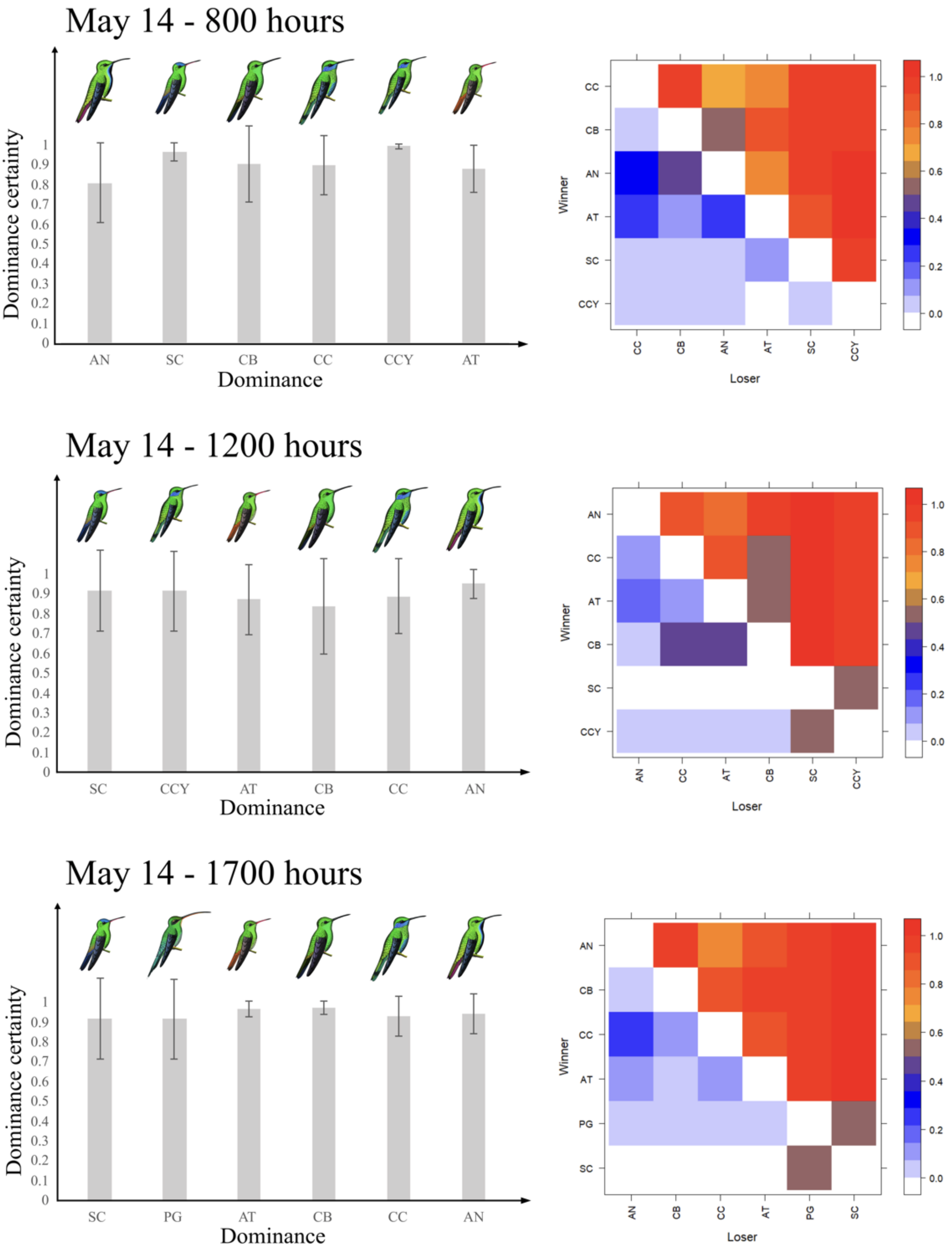


**Figure S21. Dominance certainty across hours on May 14.** Species codes are listed in Table 1. Dominance certainty was lowest at 1200 hours, while at 800 and 1700 hours, the most dominant species exhibited higher certainty in their hierarchical positions.

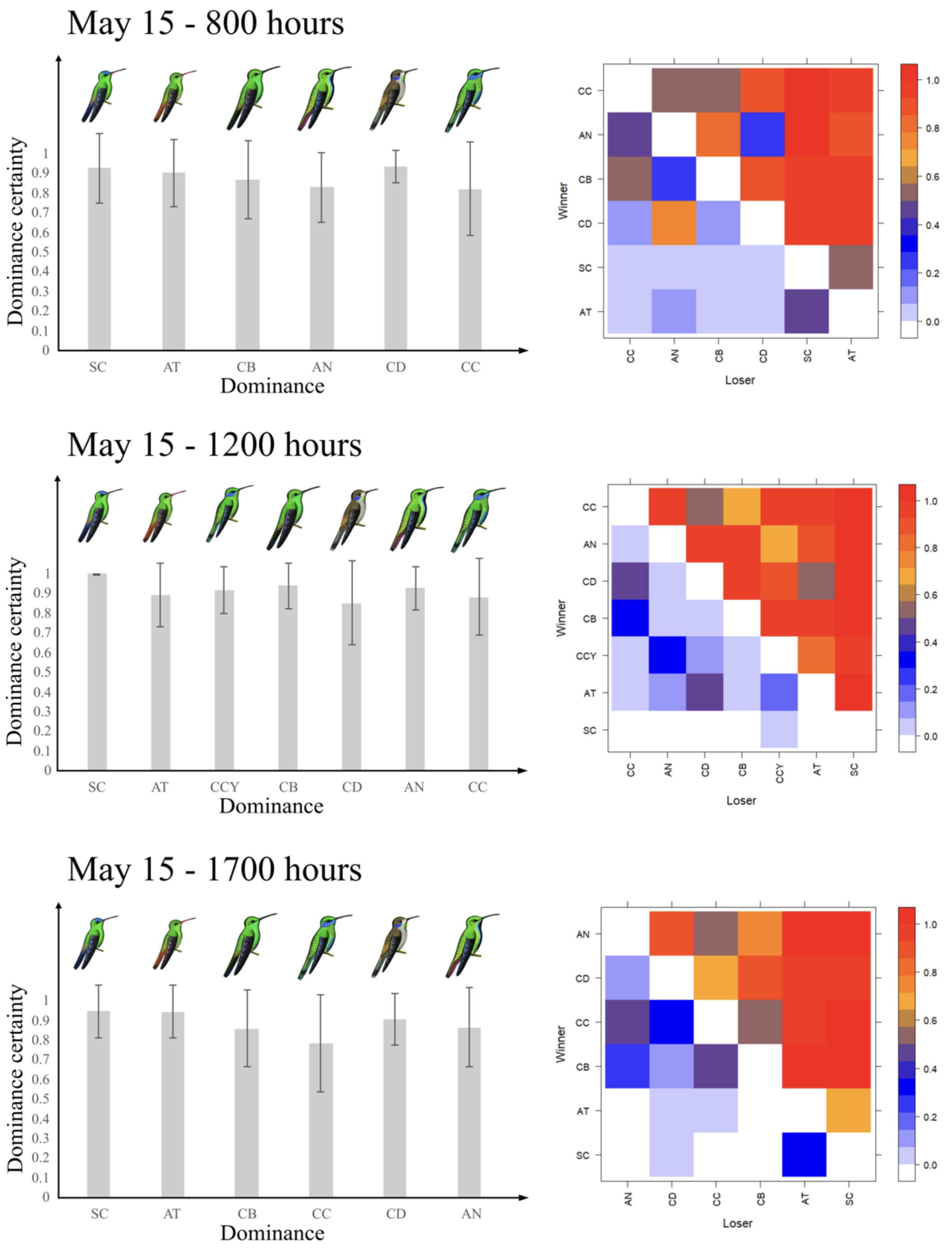


**Figure S22. Dominance certainty across hours on May 15.** Species codes are listed in Table 1. Certainty levels varied considerably throughout the day, with 1200 hours showing the highest and most stable dominance certainty among species.

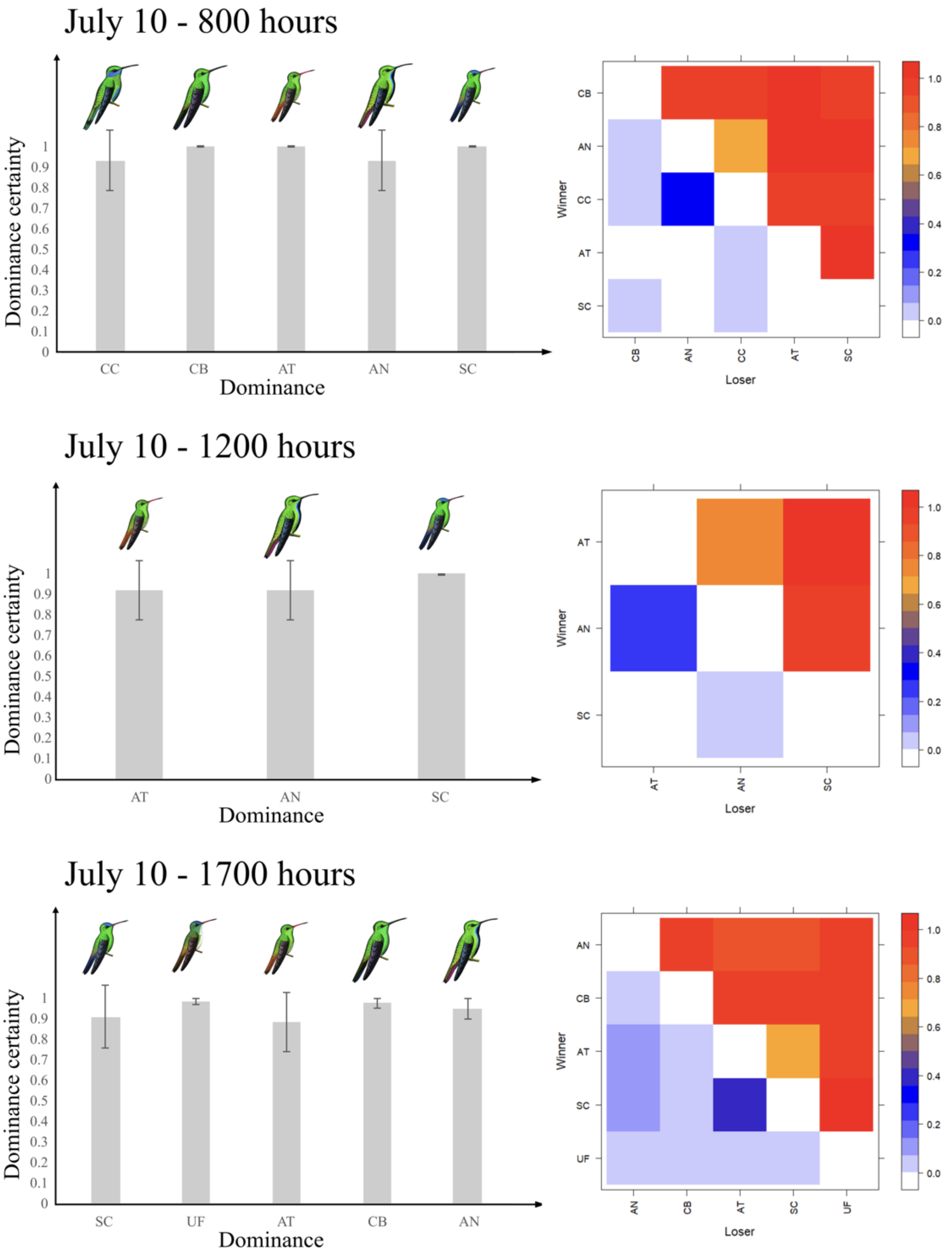


**Figure S23. Dominance certainty across hours on July 10.** Species codes are listed in Table 1. Certainty levels were highest at 800 hours but declined throughout the day. Dominant species consistently exhibited higher certainty compared to subordinates.

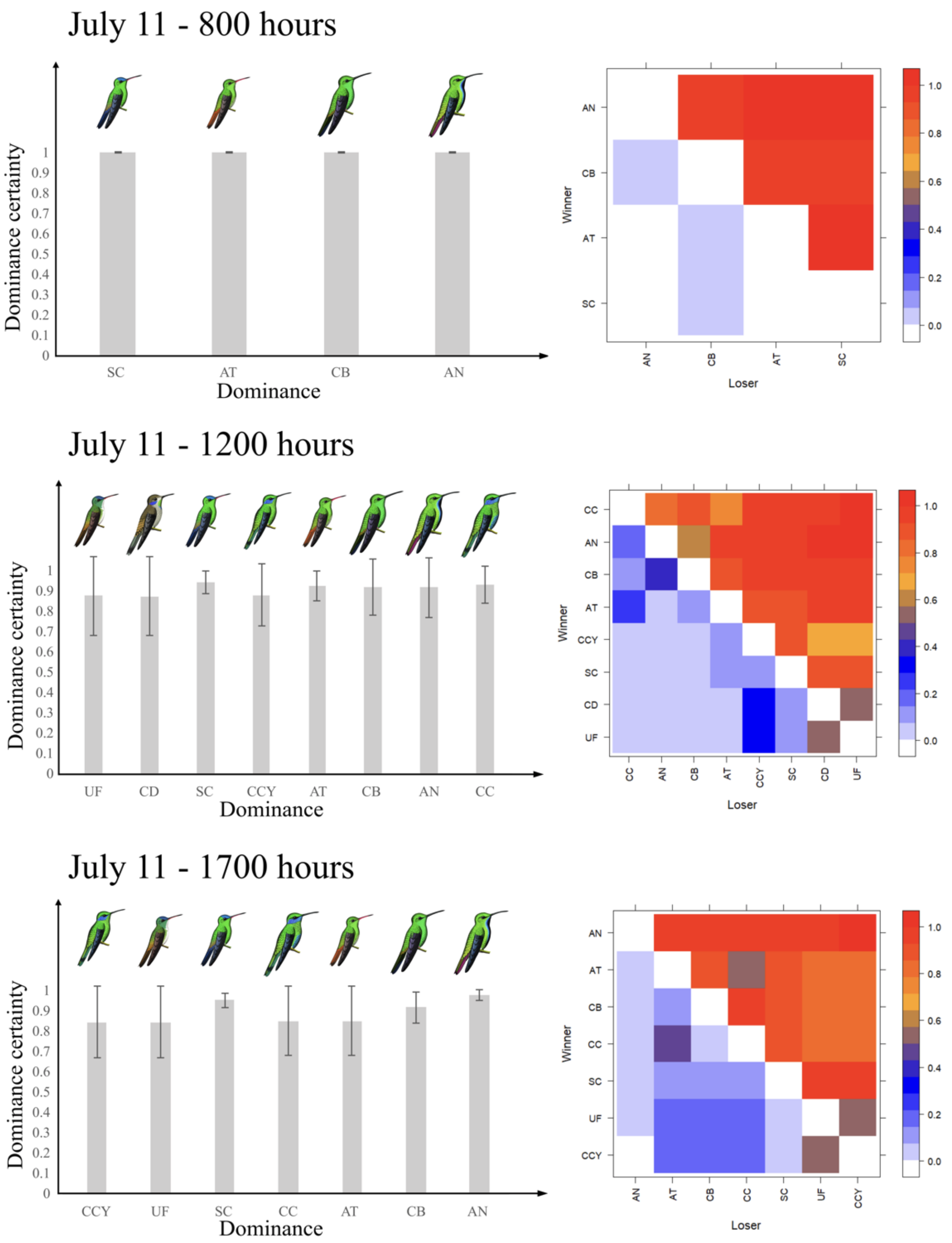


**Figure S24. Dominance certainty across hours on July 11.** Species codes are listed in Table 1. Certainty peaked at 800 hours but declined in later intervals. Throughout the day, dominant species maintained higher certainty in their hierarchical positions than subordinate ones.

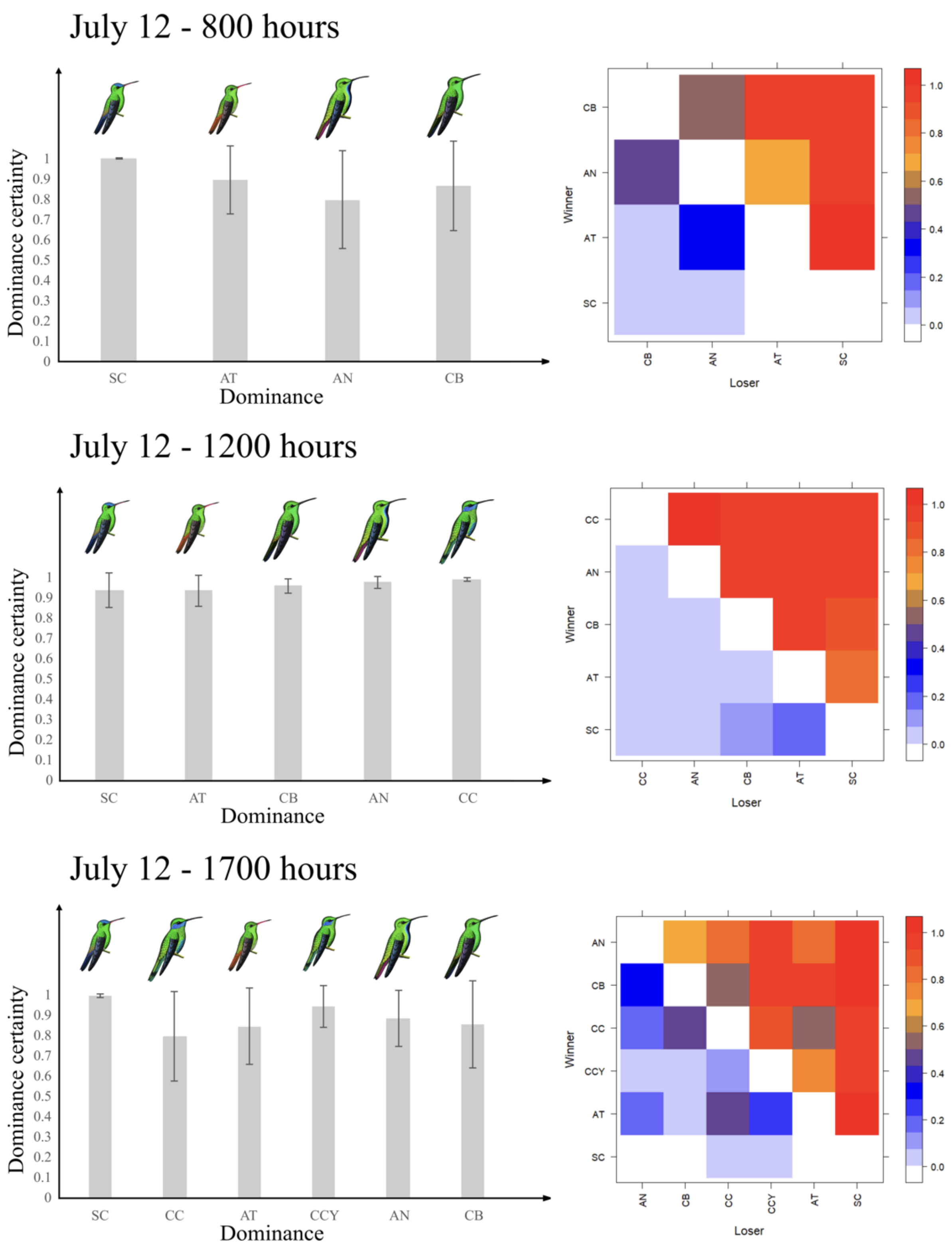


**Figure S25. Dominance certainty across hours on July 12.** Species codes are provided in Table 1. Dominance certainty fluctuated throughout the day, with the highest and most stable values observed at 1200 hours.
